# Evaluation of genotoxic and cytogenetic effects of saponins and aluminum hydroxide adjuvant on viral vaccine

**DOI:** 10.1101/2021.05.04.442590

**Authors:** Roberta Fiusa Magnelli, Rita de Cássia Stocco, Pedro Ismael da Silva Junior, Ronaldo Zucatelli Mendonça

## Abstract

Bovine papillomavirus (BPV) is the etiological agent of bovine papillomatosis, infectious disease characterized by the presence of benign tumors that can progress to malignancy. The phylogenetic classification of the PVs is performed based on the sequence homology of the Open Reading Frame L1, the most conserved among different viral serotypes. Given the immunogenicity of saponins, it,s has been used as a candidate as adjuvant use. For this reason, the safety of using saponin as an adjuvant has to be better determined to human or veterinary vaccine use. So, this study aimed to evaluate the mutagenic and genotoxic effect of saponins in comparison with the adjuvant widely used aluminum hydroxide using an isolated and purified L1 protein from BPV as model. In this study, genomic lesions, which after processed without repair can result in mutations, were detected by comet assay. Possible damages to genetic material caused by structural chromosomal changes (clastogenesis), as well as chromosomal losses (aneugenesis) were evaluated by the micronucleus test. Both tests were done on polychromatic erythrocytes and Vero cells. The evaluation of apoptosis and necrosis of treated Vero cells was made by Annexin V / PI staining and flow cytometry. The two vaccine products (L1 +Saponin and L1 + Aluminum Hydroxide) showed damages compatible with the positive control in the comet assay and both slightly elevated the micronucleus levels, in the Cell Viability Assay the results with Aluminum Hydroxide were satisfactory, characterizing Aluminum Hydroxide as a safer adjuvant according to the proposed tests, better than the saponins. Some fractions of the saponin extract separated by High Performance Liquid Chromatography were evaluated against genotoxic activity by comet assay, and their identities were confirmed by similarity to the reference standard by mass spectrometry.

## 1. Introduction

Papilloma viruses (PVs) are small circular double-stranded DNA viruses, non-enveloped and with icosahedral symmetry (DOORBAR, 2005). PVs belong to the Papillomaviridae family, being one of the oldest and most extensive virus families known (MUNDAY et al., 2014). PVs present tropism for epithelial and mucous cells, causing benign lesions (papillomas), which can spontaneously regress or progress to malignancy (STOCCO DOS SANTOS et al., 1998) In this context, BPV stands out as the best model for studying the oncogenic mechanisms of PVs in view of their genomic and pathological similarities with HPV (MUNDAY, 2014). BPV is the etiologic agent of bovine papillomatosis, an infectious and contagious disease that mainly affects young cattle. Gene products of the virus are associated with the induction and promotion of carcinogenesis. BORZACCHIELLO and ROPERTO, 2008, in a review, summarise BPV genome organization, and describe in greater detail the functions of viral oncoproteins, the interaction between the virus and co-carcinogens in tumour development and relevant aspects of immunity and vaccines. Recently, RUSSO et al, 2020, has demonstrated that BPV-2 can infect the amnion of water buffaloes and suggest that this infection may cause proliferation of the epithelial cells of the amnion.

Prophylactic vaccines against HPV have been available on the market since 2006 (RIBEIRO-MULLER, 2014). Such vaccines are based on VLPs (virus like particles) of the structural protein L1, since this can self-organize into pentamers (MARIGLIANI et al., 2012). Although two prophylactic HPV vaccines are available on the market, there are currently no commercial vaccines against BPV. The idea of developing a vaccine against papillomas started with the injection of papillomas extract in the 1940s (SHOPE, 1937). Since then, different vaccine models have been proposed in the literature (CAMPO, 2006). However, none are available on the market. Studies show that early BPV proteins (E6 and E7) have therapeutic action, while structural ones (L1 and L2), prophylactic (CAMPO et al., 1997). Studies based on structural proteins indicate that the L2 protein has a lower immunogenicity than L1, and is not a good candidate for the vaccine (RIBEIRO-MULLER, 2014). Furthermore, peptides isolated from the L2 amino-terminal portion of BPV-4 were not able to induce protection against infection by the virus (CAMPO et al., 1997). For this, adjuvants are used to enhance the response. The adjuvant must be able to promote a high and prolonged immune response and to induce a biologically active response through the modulation of the immune system (AUDIBERT, 2003), in addition to directing that immune response to a protective response, preventing the disease (MOREIN et al., 1996).

Natural bioactive compounds derived from plants and the marine environment are well known for their pharmacological properties and therapeutic effects in the prevention and treatment of various diseases. In this sense, saponins present themselves as potent candidates for adjuvants in vaccines based on VLPs (PENG et al., 2015).

However, before developing pharmaceutical agents from natural sources, it is important to research them for their cytotoxicity and hemolytic activity in healthy erythrocytes (KUMAR et al., 2011, SHARMA et al., 2001). Saponins, an important class of natural products, have a wide spectrum of biological factors and pharmacological activities (CHWALEK et al., 2006, GAUTHIER et al., 2009). These produce foam when mixed with water. The amphiphilic properties of saponins allow them to interact with cell membranes (SOLTANI et al., 2014, AMINI et al., 2014). Although the mechanisms involved in hemolysis, promoted by saponins, have not been studied, saponins are known to interact with cholesterol in erythrocyte membranes, forming pores that destabilize them (CHWALEK et al., 2006, GAUTHIER et al., 2009) . This activity leads to the release of hemoglobin and other components in the surrounding fluids.

Many studies have been carried out exposing the high immunogenic capacity and low hemolytic activity of saponins, but none show the structure-activity relationship for this general class of compounds, which is due to the lack of systematization of available information (ORTEGA et al., 2009). The relationship between adjuvant and hemolytic activity has already been reported, but these studies were based on a few samples (KENSIL et al., 1991; BOMFORD et al., 1992).

However, natural products are not intrinsically safe and effective, and can cause damage to health (NOHYNEK et al., 2010), which justifies the need to assess the safety of new plant raw materials, thus guaranteeing good condition of use (ANTIGNAC et al., 2011).

For human vaccines, registered adjuvants are limited and include aluminum and oil / water based adjuvants. These adjuvants induce robust antibody responses, but weak cell-mediated immunity. Saponin-based adjuvants (SBAs) increase the Th1 response by stimulating cytokines (IL-2 and INF-γ) (SINGH AND O’HAGAN, 2003, KENSIL, 1995).

Currently, other adjuvants have been studied, some based on plant extracts. The Agave sisalana Perrine (Figure 4), popularly known as sisal, it is a herbaceous originated in Mexico and adapted to the semi-arid climate of the Caatinga (GUTIERREZ et al., 2008). This species is distributed in the Northeast of Brazil, where its cultivation occupies extensive areas of poor soils, being considered an important productive alternative for this region (RIBEIRO FILHO, 1967), playing an important socioeconomic role, especially to family farmers, since the management of tillage, harvesting, defibration, fiber processing, industrialization and / or making of handicrafts (MARTIN, 2009).

The spirostane saponins found in Agave are an important class of natural products for the pharmaceutical industry because they are raw materials for the production of sex hormones (mainly contraceptives, corticosteroids, steroidal diuretics and vitamin D) (HOSTETTMANN; MARSTON, 1995). Studies of adjuvant and hemothilic activity in *Agave sisalana* are still scarce, as well as safety in its pharmaceutical use.

### Comet Assay as a genotoxicity indicator

Genotoxicity is a recent specialty, and is located at the interface between toxicology and genetics, which is why it is often called toxicological genetics. This aims to study the processes that alter the genetic basis of life, whether in its physical-chemical structure, deoxyribonucleic acid (DNA), a process classified as mutagenesis; whether in altering genetic determinism at cellular and organic levels, identified, respectively, as carcinogenesis and teratogenesis (SILVA et al. 2003).

The comet assay (EC) was developed by Östling and Johanson in 1984 and modified by Singh et al. (1988), who attributed greater sensitivity to the technique with the use of alkaline solution, being considered effective in cytogenetic tests (FLORES; YAMAGUCHI, 2008).

The EC is used to detect DNA damage, being the main integrator and complementary to other mutagenic tests (TICE et al., 2000) with the purpose of identifying breaks induced by mutagens in the DNA molecule (FABIANI et al., 2008).

EC is not used to detect mutations, but DNA damage that, after being processed, can result in mutations. Unlike the mutations, the lesions detected by the test are amenable to correction, so the test needs very well-established controls as it is very sensitive, the time between exposure and analysis must be as short as possible, as the damage can be repaired, so the short detection time. Therefore, the comet test can also be used for DNA repair studies, (ALBERTINI et al. 2000).

The Butantan Institute has been dedicated to the study of BPV for more than two decades, contributing to different research segments, such as the discovery of new viral types in the State of São Paulo, new diagnostic methods, viral isolation, expression of recombinant proteins, analysis of cytogenetic changes and pathological studies of papillomas and carcinomas associated with the virus Bovine papillomatosis is not only a problem of buiatria, but also an economic one, as this branch represents one of the main segments of Brazilian agribusiness, covering two highly profitable aspects: the meat and milk production chains.

Furthermore, BPV is considered the best study model for HPV. There is a wide variety of studies on prophylactic and therapeutic vaccines against human papillomavirus (HPV), which is of great relevance in the area of human health due to its association with various types of malignancies, including cervical and head and neck cancer.

The procedures developed have been the starting point for studies on vaccine strategies, and obtaining recombinant proteins and saponins in the laboratory allows a detailed study of their effects. Cloning in bacterial vectors for the expression and purification of proteins has advantages such as the large amount of biomass generated, low relative cost and speed, serving different purposes, however, in order to arrive at a final product, many steps must be taken. For this reason, this study is an extremely important proposal as a pre-clinical trial.

Therefore, this work aims to evaluate the mutagenic and cytotoxic potential of adjuvant as saponin and aluminum hydroxide using BPV L1 protein associated as model. A precise knowledge of the safety of the use of these adjuvants is of great importance, especially with the current development of new vaccines against the pandemic of COVID-19

## 2. MATERIALS AND METHODS

### 2.1. Cells

Monolayer cell cultures of Vero cells (African green monkey kidney-ATCC CCL-81), with passages between 148-152, were grown at 37°C and 5% CO 2 in minimum essential medium (MEM) supplemented with 4 mM glutamine, and 10% fetal bovine serum using standard cell culture techniques. Viable cell counts were performed in Neubauer chamber using Trypan blue (0.05%) exclusion.

### 2.2. Blood Collection and DNA Extraction

#### 2.2.1. Sampling

10 mL of peripheral blood were collected from ten cattle (*Bos taurus*, Simental) by venipuncture of the jugular vein, using vacuum tubes containing potassium heparin. Blood samples were collected in the city of Itapetininga and transported to the Genetics Laboratory of the Butantan Institute in thermal boxes to prevent possible damage to the genetic material, as well as to preserve viable cells, thus allowing the incubation of biological material for the accomplishment of the Micronucleus test with cytokinesis block performed on the same day.

#### 2.2.2. Extraction of genomic and viral DNA from collected blood

The material used for DNA extraction was kept at -20ºC. The DNA was extracted using the QIAamp® DNA Blood Mini Kit (Qiagen, Hilden, Germany), according to the manufacturer’s instructions. The extracted DNA was quantified in a BioPhotometer Plus spectrophotometer (Eppendorf, Hamburg, Germany), where a 2 μL aliquot of the respective samples was read at wavelengths of 230, 260 and 280 nm, calculating the concentration in ng / μL, by multiplying the absorbance value by the extinction coefficient.

### 2.3. Molecular BPV Identification and Typing

The blood collected was subjected to molecular identification of BPV sequences using the set of primer pairs shown in Table 1.

**Table 1.**
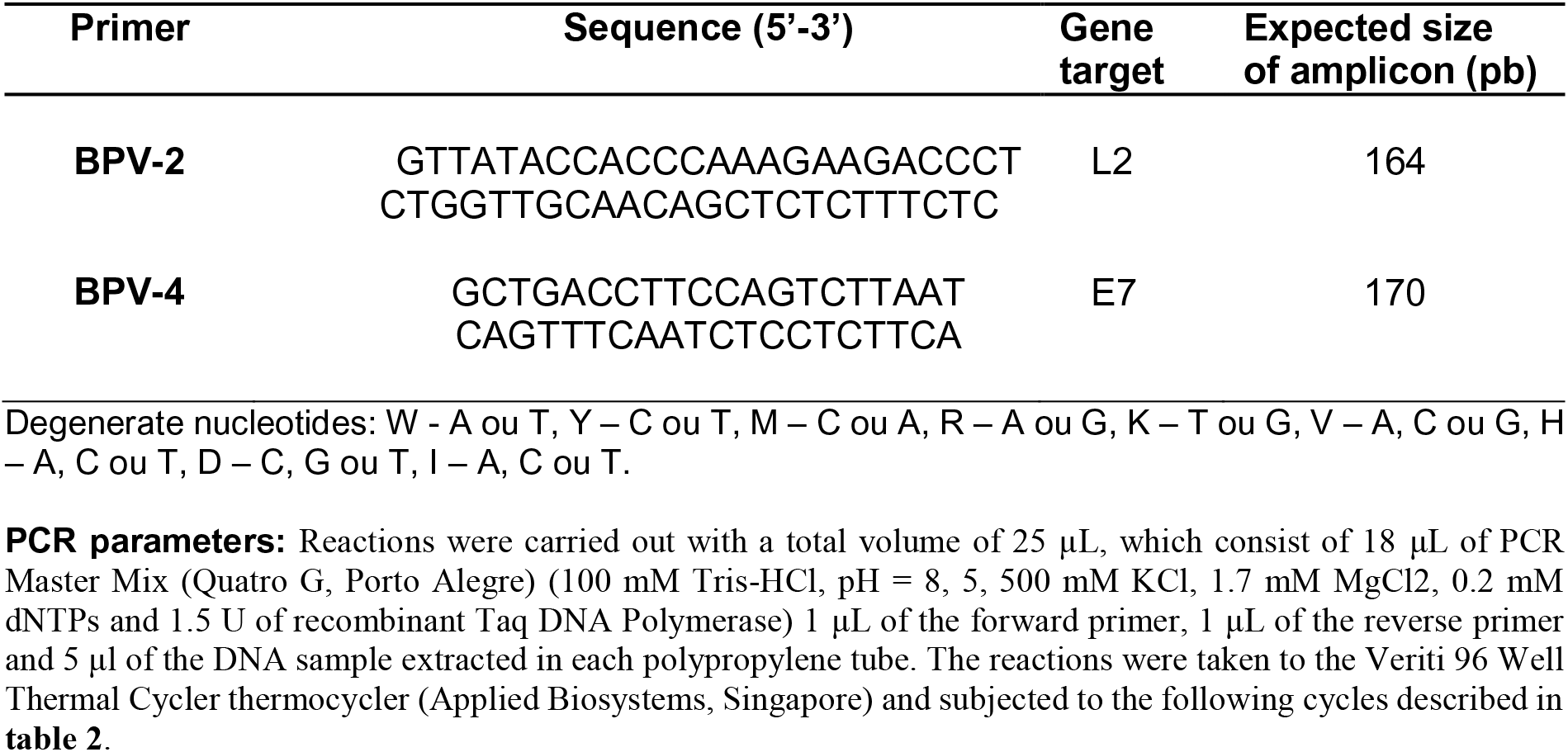
Sequences of primers for diagnosis.

In table 2 is showed the PCR reaction parameters used

**Table 2.**
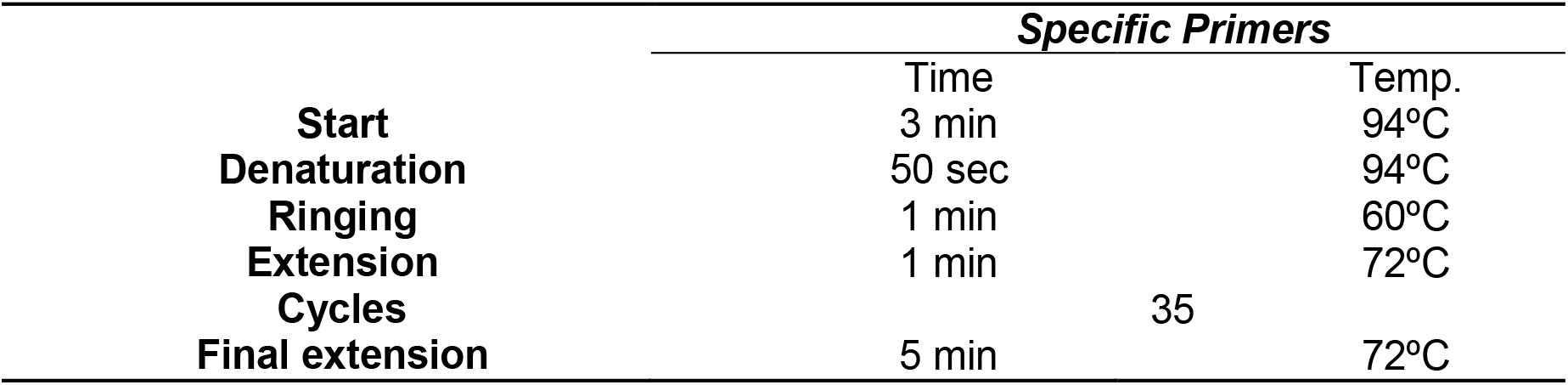
PCR reaction parameters using the different pairs of primers.

### 2.4. Obtaining sisal residue and preparing extracts

#### 2.4.1. Obtaining sisal juice

The raw material extracted in Bahia (Sisal juice) obtained directly from the defibration process of *Agave sisalana* leaves in a sisal cultivation farm located in the city of Valente, Bahia (latitude 11 ° 44 ′ 24 ″ S, longitude 39 ° 271 43 ″ W and altitude 585m) through a partnership with the Secretariat of Science, Technology and Innovation of the State of Bahia (SECTI) and Laboratory of Pharmaceutical Technology in Phytotherapy on the UNESP-Assis campus. The leaves were dried at 50ºC using an electric dryer and crushed with the aid of a mechanical shredder until it became a powder.

#### 2.4.2. Obtaining the dry sisal precipitate

To obtain the dry precipitate of A. sisalana (DPAS), the sisal juice was centrifuged (FANEM FR 22) at 448G for 20 minutes with a centripetal force of approximately 52,580 N. Subsequently, the supernatant was dried in an oven (FANEM LTDA, model 002CB) at 40 ° C, until constant weight is obtained.

#### 2.4.3. Obtaining the extract by acid hydrolysis

To obtain the acid hydrolysis extract of *A. sisalana* (AHEAS), the sisal juice was heated to 100ºC ten times and hydrolyzed with 2N HCL for four hours, under agitation. The precipitate was separated from the acidic solution by filtration at room temperature.

The extracts were filtered, lyophilized and kept at 4ºC until use. This processing and the study concentration were obtained by the Biotechnology laboratory, at Julio Mesquita University (UNESP-Assis) and forwarded to the Butantan Institute for collaboration in carrying out this work.

### 2.5. L1 Protein Obtaining

The BP1-L1 gene, previously cloned into pAT153, was inserted into a pET28a expression vector. The vector called pET L1 was cloned into dH5 bacteria and subsequently inserted into BL21 bacteria for protein expression. E. coli modified BL21 (DE) were used for protein expression. Recombinant protein expression was evaluated by Western Blot and mass spectrometry. Protein purification was performed by affinity chromatography and ion exchange, then undergoing spectroscopic analysis by circular dichroism to assess its folding and thermal stability. The L1 protein was expressed and purified in the Genetics laboratory of the Instituto Butantan, by Modolo (2014), and assigned to this work for the mutagenicity tests in the concentration of 1ug / mL according to Modolo et al, 2017 .

### 2.6. Micronucleus Test with Cytokinesis Block (TMBC) in Peripheral Blood

The collected blood was incubated in 15 mL tubes containing: 4.5 mL of RPMI 1640 medium, supplemented with 0.5 mL of fetal bovine serum and 0.1 mL of phytohemagglutinin A (PHA). 200 µL of whole blood was added to each tube. The incubation of whole blood is preferably recommended, as the isolation of peripheral blood mononuclear cells with Ficoll-Paque can induce DNA damage (ARALDI et al.,2015) The tubes were incubated in an oven at 37ºC.

After 8 h of incubation the 10 samples were treated according to table 3:

**Table 3:**
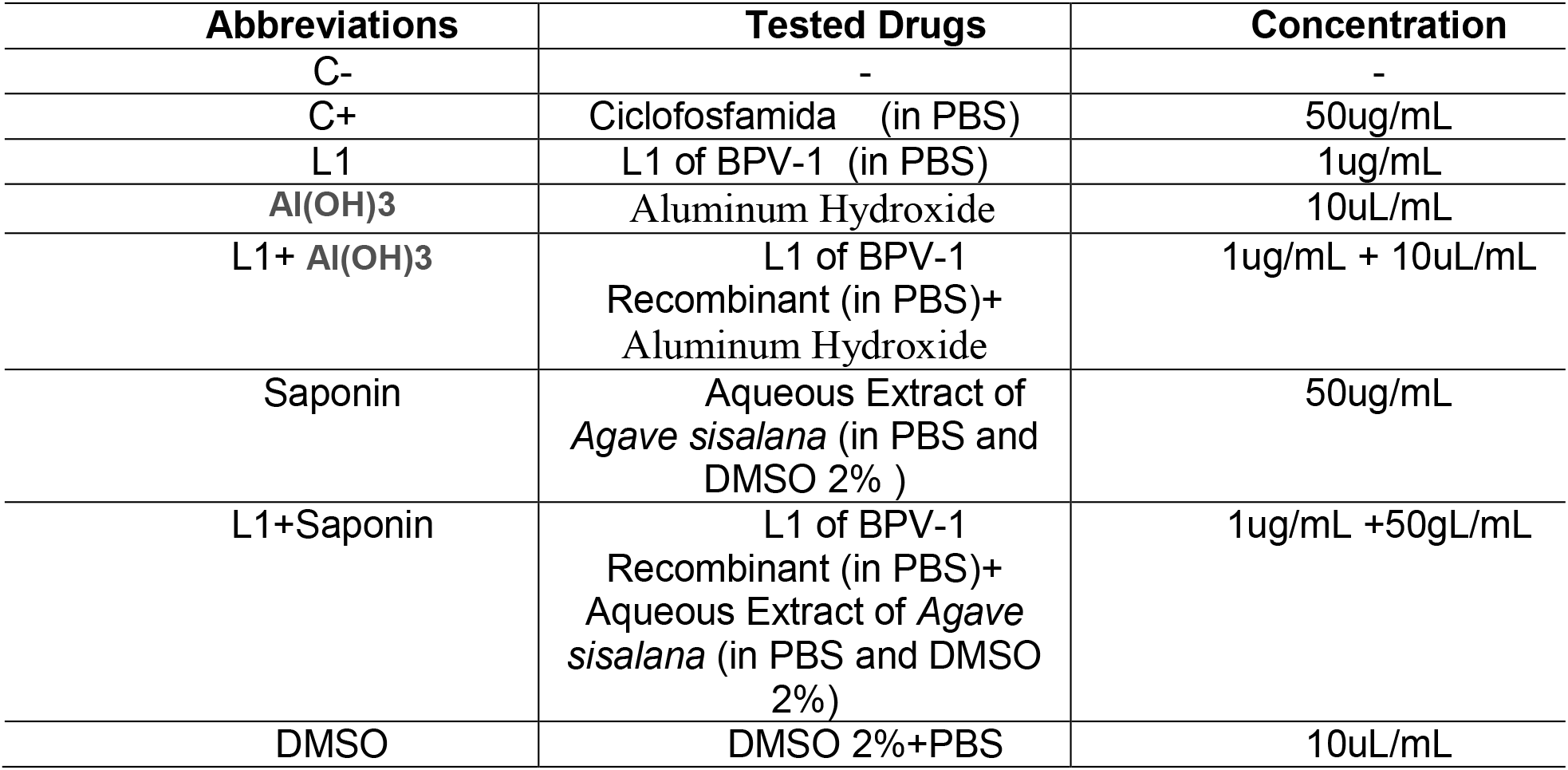
Drugs tested and work concentrations

After 44 h of the beginning of the procedure, the collected samples were treated with 6 µL of cytochalasin B, diluted in DMSO, in the concentration of 6 µg / mL, to prevent cytokinesis, according to Araldi et al (2015). The samples were incubated for 72 h and then 0.5 ml of Carnoy fixative (3: 1 methanol-acetic acid) was added for 5 min at room temperature. The tubes were centrifuged for 10 min at 112 xg. The supernatant was discarded by inversion. 5 ml of Carnoy fixative was added per tube. The material was carefully homogenized with the aid of disposable Pasteur pipettes to avoid possible contamination. The tubes were centrifuged for 10 minutes at 112 xg, discarding the supernatant. The procedure was repeated three times.

After the last centrifugation, the material was aspirated and transferred to slides previously cleaned with 70º alcohol. The slides were dried at room temperature overnight, fixed in absolute methanol and stained with Giemsa 2% in PBS buffer pH 6, 8 for 8 min. The material was washed three times in distilled water for 3 min. The slides were mounted with Entelan (Merck, Germany) and analyzed using a Primo Star binocular light microscope (Zeiss, Germany) with a 100 X objective.

1,000 cells per slide were analyzed, observing the frequency of micronucleated lymphocytes. The analysis of the TMN in peripheral sage is performed by evaluating the frequency of micronucleus formation (MNr0 = a / b, where: a - number of lymphocytes with MN and b - total number of lymphocytes). Based on the number of micronucleated cells, a histogram is performed using Excel software.

### 2.7. Micronucleus Test with Cytokinesis Block in Vero Cell Culture

TMNBC was also carried out on Vero epithelial cell lines not infected by BPV as described by FLORES and YAMAGUCHI, 2008.

Cell culture: Eight cultures treated according to table 3 were established. Substances were added and after 1 h 6 µg / mL of Cyt-B were added. The material was incubated for 48 hours, since the cell duplication time for the strain used is 24 hours, thus ensuring two replication cycles.

After incubation, the medium was removed with a sterile pipette. The plates were washed with 2 ml of PBS at 37ºC, which was discarded by inversion. 2 ml of trypsin-EDTA solution was added at 37ºC. The plates were incubated for 5 minutes. After this time, the material was aspirated and transferred to the Falcon tube containing the medium initially removed, allowing inactivation of trypsin. The material was centrifuged for 5 minutes at 112 xg. The cell pellet was collected and transferred to previously washed slides, using a smear. The blades were assembled using Entellan (Merck, Germany). The material was analyzed using a Primo Star binocular microscope (Zeiss, Germany) with a 100 X objective.

1,000 cells per slide were analyzed. Analyzes of TMN in Vero cells were performed by evaluating the frequency of micronucleus formation (MNr0 = a / b, where: a - number of cells with MN and b - total number of cells). Based on the number of micronucleated cells, a histogram was performed using Excel software.

### 2.8. Comet assay (CA) in blood cells

CA was used to analyze the mutagenic potential in blood cells.

#### Pre-preparation of the slides

26 x 76 mm slides were immersed in a beaker containing normal melting point agarose solution (NMA - normal melting agarose), diluted in PBS buffer free of Ca2 + and Mg2 + at 1.5% at 60ºC, having one side cleaned with paper towels. The slides remained overnight at room temperature, in a horizontal position, for the complete drying of the agarose.

#### Sample treatment

The blood material collected from the ten calves was divided into eight samples of 200 μL each. The aliquots were distributed in 1.5 ml polypropylene tubes containing 200 μL of RPMI 1640 medium and the drugs were applied to the tubes as shown in table 3. The material was incubated for one hour in an oven at 37º C, according to Araldi et al . (2015) [39] . After this period, the tubes were centrifuged at 1120 xg for one minute, discarding the supernatant. A volume of 10 μL of the centrifuged material was transferred to polypropylene tubes of 0.2 mL and added 75 μL of low melting agarose (LMA) at 37º C. The final volume of 85 µL of the suspension cell was aspirated with the aid of a pipette and transferred to slides pre-covered with 1.5% NMA.

The slides were covered with a coverslip and incubated at 4 ° C for 20 minutes for the agarose to solidify. After this time, the coverslip was carefully removed. The slides were inserted into a flask containing cold lysis solution (71 mL of the stock solution of 2.5 M NaCl, 100 mM EDTA, 10 mM Tris-HCl, plus 0.8 mL of Triton X-100 and 8 mL of DMSO), remaining in it for one hour at 4º C. From this moment on, the remaining steps were performed with the laboratory lights off to avoid any damage to the DNA.

#### Electrophoresis

After lysis, the slides were washed in PBS for five minutes. The material was transferred to a horizontal vat, and the electrophoresis buffer (300 mM NaOH, 1 mM EDTA, pH> 13.0) was added. The material remained immersed in the buffer for 40 minutes to remove histones. After this period, the electrophoretic run was started under the following conditions: 24 V (0.74 V / cm), 300 mA for 30 minutes. The electrophoretic vat was coated with ice sheets to ensure that the material remained at 4º C during the experiment.

#### Neutralization

After electrophoresis, the material was neutralized in a neutralizing solution (400 mM Tris-HCl, pH 7, 5) for five minutes.

#### Material analysis

The material was stained with 20 µL of propidium iodide (PI) 4 µg / mL and analyzed using an Axio Scope A1 epifluorescence microscope (Carl Zeiss, Germany) at a total magnification of 400 X. 100 nucleoids per slide were analyzed, in a total of 8 slides (800 nucleoids) for each of the 8 samples, totaling 6,400 nucleoids, which were classified into: 0 (no damage), 1 (intermediate damage) and 2 (maximum damage), as shown in figure 6.

Statistical analysis: Based on the number of nucleoids observed per class, the score value per slide was obtained, according to the formula:

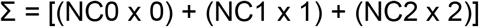

Where: NC0 (number of nucleoids observed in class 0), NC1 (number of nucleoids observed in class 1) and NC2 (number of nucleoids observed in class 2).

Based on the obtained score values, the Kruskal-Wallis test was performed followed by the Dunn post-hoc test, both with a 5% significance level using the GraphPad Prism software version 5 (GraphPad Software, Inc.).

## 2.9. Comet assay (CA) in Vero cell culture

Eight culture flasks of 25 cm2, with confluence of 80% of VERO cells were destined for the comet assay. Drugs were applied to the flasks according to table 3. The material was incubated in an oven at 37ºC, with an atmosphere of 5% CO2 until reaching 100% confluence. The cells were washed with 2.0 ml of sterile PBS and centrifuged at 1120G, discarding the supernatant. The pellet was homogenized with 200 μL of PBS. A volume of 20 μL of the cell suspension was transferred to a polypropylene tube of 0.2 mL and added 170 μL of low melting point (LMA) agarose (Fermentas, Lithuania) at 0.8%, diluted in PBS at 37ºC. The material was homogenized and transferred to the slides pre-covered with 1.5% NMA agarose and covered with a coverslip.

The slides were covered with a coverslip and incubated at 4 ° C for 20 minutes for the agarose to solidify. After this time, the coverslip was carefully removed. The slides were inserted into a coply containing cold lysis solution (71 mL of the stock solution of 2.5 M NaCl, 100 mM EDTA, 10 mM Tris-HCl, plus 0.8 mL of Triton X-100 and 8 mL of DMSO), remaining in it for one hour at 4º C. From this moment on, the remaining steps were performed with the laboratory lights off to avoid any damage to the DNA. Electrophoresis and neutralization were performed similarly to the blood cell assay.

### Material analysis

The material was stained with 20 µL of propidium iodide (PI) 4 µg / mL and analyzed using an Axio Scope A1 epifluorescence microscope (Carl Zeiss, Germany) at a total magnification of 400 X. 50 nucleoids per slide were analyzed, in the total of 3 slides (150 nucleoids) for each of the 8 samples, totaling 1,200 nucleoids which were classified as: 0 (no damage), 1 (intermediate damage) and 2 (maximum damage), as shown in figure 6.

### Statistical analysis

Based on the number of nucleoids observed per class, the score value per slide was obtained. Based on the obtained score values, the Kruskal-Wallis test was performed followed by the Dunn post-hoc test, both with a 5% significance level using the GraphPad Prism software version 5 (GraphPad Software, Inc.).

## 2.10. Flow cytometry: test with Annexin V-PI

Eight culture flasks of 25 cm2, containing 5mL of minimal medium (MEM supplemented with 10% fetal bovine serum) with confluence of 80% of VERO cells were destined for the Flow Cytometry assay.

The cells were incubated with different drugs tested (table 3) at 37ºC for 48 hours, the time required for two duplication cycles. Then, the medium was transferred to 15.0 ml Falcon tubes and the cells were washed with 2.0 mL of sterile PBS and incubated with 2.0 mL of trypsin / EDTA solution (Cultilab, Campinas, Brazil) at 37ºC for five minutes to promote the monolayer breakdown. The cell suspension was transferred to the Falcon tube containing the removed medium and centrifuged at 157 xg for five minutes, discarding the supernatant. The pellet was homogenized with 1.0 mL of cold sterile PBS and transferred to a 1.5 mL polypropylene tube. The cells were centrifuged at 157 xg for five minutes, discarding the supernatant. The cells were homogenized with 100 μL of binding buffer and incubated with 5.0 μL of Annexin V-FITC and 5.0 μL of propidium iodide (PI) for 15 minutes at room temperature. The cells were homogenized with 100 μL and then centrifuged at 157 xg for five minutes, discarding the supernatant. The cells were homogenized with 100 μL of cold binding buffer and analyzed in flow cytometry of BD Accuri C6 (BD Biosciences, USA), using channels FL1 (Annexin V-FITC) and FL3 (PI). The analyzes were performed in triplicate, with 10,000 events being analyzed. The data were analyzed using the BD Accuri C6 software. Statistical analysis was performed based on the average percentage of living cells, using the two-way ANOVA test followed by the Tukey post-hoc test (P <0.05) using the GraphPad Prism software version 5 (GraphPad Software, Inc.).

## 2.11. *Agave sisalana* Perrine crude extract: chromatographic fractionation

52.3 mg of the hydrolyzed extract and also of the Saponin Sigma standard were weighed, then homogenized in 2 mL of aqueous solution of trifluoroacetic acid-0.05% TFA and subjected to High Performance Liquid Chromatography on column C18, UFLC system Shimadzu model Proeminence. The column was previously equilibrated with solvent A (TFA 0.05%) and elutions were performed in a linear gradient 0-80% acidified acetonitrile (ACN / TFA) for 60 minutes, with a flow rate of 2 mL / min. The separation was carried out at constant temperature and the absorbance was monitored at 225 and 280 nm.

The fractions corresponding to the peaks were collected manually and concentrated in a vacuum centrifuge (Savant Instrument Inc.), lyophilized (Lyophilizer Epsilon 2_6D LSC, Christ, Germany) and stored at -80ºC until the time of the tests.

## 2.12. Comet assay (CA) in Vero cell culture of the fractions obtained

The fractions chosen for analysis of genotoxic activity were those whose peaks eluted at similar retention times, comparing sample and standard. This analysis was carried out in order to discover, isolate and identify the substance that, possibly, confers genotoxicity in the composition of this compound.

Sixteen 25 cm2 culture flasks, containing 5 mL of minimal medium (MEM supplemented with serum fetal bovine 10%) with confluence of 80% of VERO cells were destined for the comet assay. The fractions obtained were reconstituted in 300 μL of DMSO 2%, and 10 μL were applied to the flasks, as well as the positive control Cyclophosphamide 50μL / mL and the diluent DMSO 2%. The material was incubated in an oven at 37ºC, with an atmosphere of 5% CO2 until reaching 100% confluence. The cells were washed with 2.0 mL of sterile PBS and centrifuged at 1120 xg, discarding the supernatant. The pellet was homogenized with 200 μL of PBS. A volume of 20 μL of the cell suspension was transferred to a polypropylene tube of 0.2 mL and added 170 μL of low melting point (LMA) agarose (Fermentas, Lithuania) at 0.8%, diluted in PBS at 37ºC. The material was homogenized and transferred to the slides pre-covered with 1.5% NMA agarose and covered with a coverslip.

The slides were covered with a coverslip and incubated at 4 ° C for 20 minutes for the agarose to solidify. After this time, the coverslip was carefully removed. The slides were inserted into a coply containing cold lysis solution (71 mL of the stock solution of 2.5 M NaCl, 100 mM EDTA, 10 mM Tris-HCl, plus 0.8 mL of Triton X-100 and 8 mL of DMSO), remaining in it for one hour at 4º C. From this moment on, the remaining steps were performed with the laboratory lights off to avoid any damage to the DNA. Electrophoresis and neutralization were performed similarly to the blood cell assay.

### Material analysis

The material was stained with 20 µL of propidium iodide (PI) 4 µg / mL and analyzed using an Axio Scope A1 epifluorescence microscope (Carl Zeiss, Germany) at a total magnification of 400 X. 100 nucleoids per slide were analyzed, in the total of 3 slides (300 nucleoids) for each of the 16 samples, totaling 4,800 nucleoids which were classified as: 0 (no damage), 1 (intermediate damage) and 2 (maximum damage), as shown in figure 6. Based on in the number of nucleoids observed per class, the value of the score per slide was obtained. The three score values per slide were added for each sample so that an activity comparison could be made between the extract and the Sigma standard. The fractions that obtained a score value closer to the positive control were chosen for further analysis in a mass spectrometer.

## 3. RESULTS

### 3.1. DNA extraction and quantification

10 ml of peripheral blood were collected from ten cattle (Bos taurus, Simental) up to 2 months old, since studies point to a correlation of age-dependent DNA damage (HEUSER et al., 2008).

The extracted DNA was quantified in a BioPhotometer Plus spectrophotometer (Eppendorf, Hamburg, Germany), where a 2 μL aliquot of the respective samples was read at wavelengths of 230, 260 and 280 nm. The DNA concentration was calculated in ng / μL and the degree of purity in the 260/230 and 260/280 ratios according to table 4.

**Table 4.** Spectrophotometer determination of the DNA concentration of the collected blood samples and the different ratios at different wavelength.

| Samples | ng/μL | R (260/230) | R (260/280) |
| --- | --- | --- | --- |
| 1 | 17 | 1, 07 | 1, 03 |
| 2 | 27 | 0, 93 | 1, 08 |
| 3 | 27 | 0, 93 | 1, 08 |
| 4 | 65 | 1, 00 | 1, 57 |
| 5 | 70 | 0, 82 | 1, 39 |
| 6 | 65 | 1, 06 | 1, 58 |
| 7 | 43 | 0, 83 | 1, 70 |
| 8 | 37 | 1, 32 | 1, 26 |
| 9 | 80 | 1, 90 | 1, 64 |
| 10 | 26 | 3, 15 | 1, 49 |

### 3.3. Molecular BPV identification and typing in peripheral blood samples

Blood samples collected from ten calves were subjected to molecular identification of viral DNA sequences by means of PCR. For this purpose, three pairs of specific primers (BPV-1, BPV-2 and BPV-4) were used. The results of identifying BPV sequences are shown in the figure 1.

**Figure 1.**
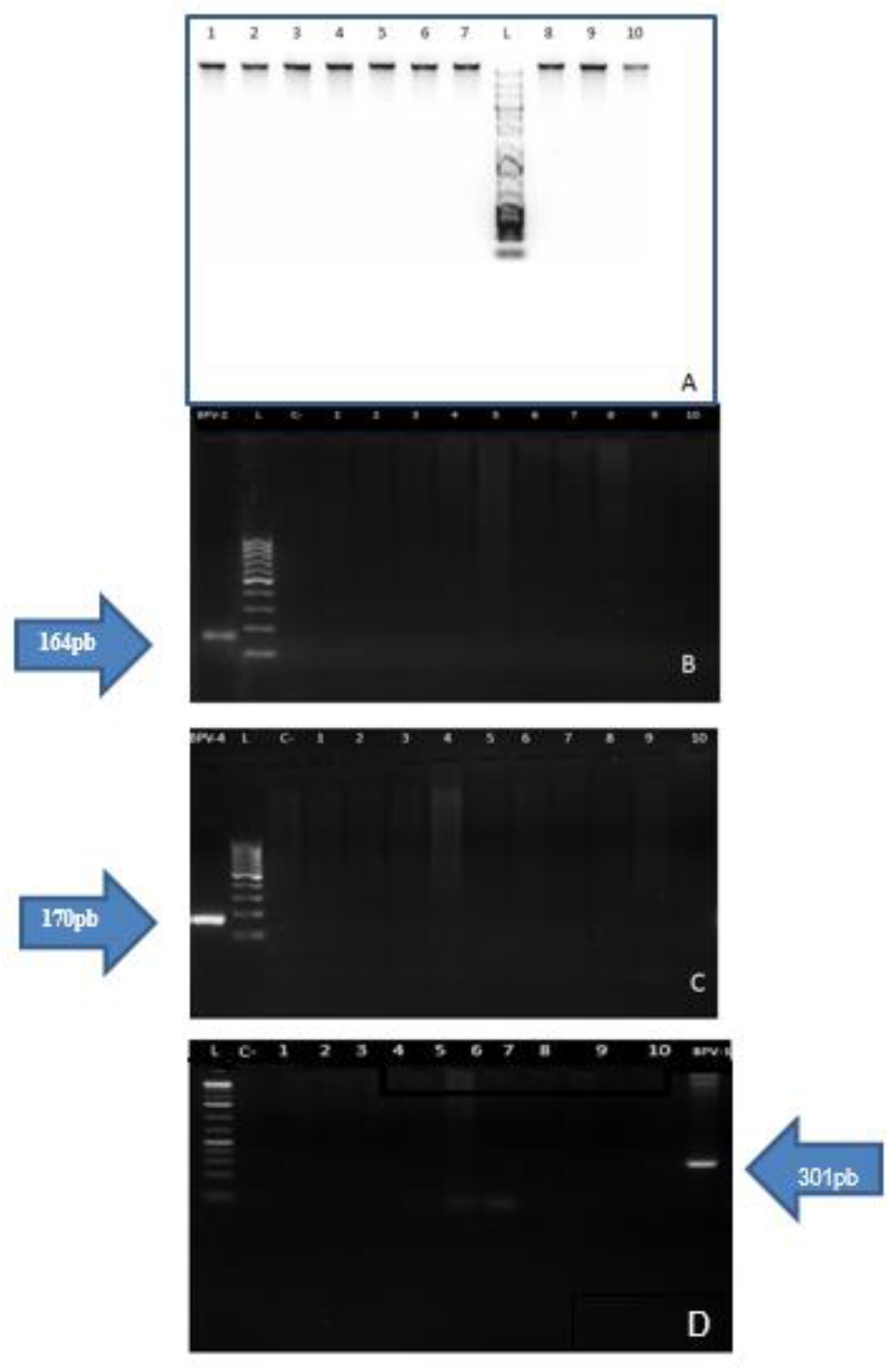
Images of electrophoresis gels showing: the integrity of the extracted genomic DNA. A 1 Kb DNA Ladder marker (Invitrogen, Carlsbad, USA) (A) was used and the absence of amplicons was verified using specific primers for BPV-2 (B), BPV-4 (C) and BPV-1 (D). The images show only the presence of amplicons consistent with the positive controls. Marker used 100 bp DNA Ladder (Invitrogen, Carlbad, USA) in figures B, C and D.

The molecular diagnosis of the ten peripheral blood samples revealed the absence of amplicons for the three different pairs of primers used (figure 7). However, the primers amplified the positive control samples made up of BPV-1, 2 and 4 genomes, cloned in *E. coli* D5Ha bacteria. No amplicons were observed in the negative controls, eliminating the possibility of contamination.

These results demonstrate the absence of previous BPV infection in the ten samples. The absence of the virus in blood samples is an important finding, given that studies have shown that cells infected by BPVs 1, 2 and 4 have high clastogenic levels (ARALDI et al., 2013, ARALDI et al., 2014). Some blood samples may present the virus, but without expression in epithelium, as they are in a latent state (SILVA et al., 2013). In asymptomatic cattle, the activation of the virus in the blood can be independent of the productive infection in the epithelial tissue (CAMPO et al., 2006). The presence of BPV in the blood of newborn calves has already been detected by other authors (STOCCO DOS SANTOS et al., 1998; FREITAS et al., 2003; YAGUIU et al., 2008), suggesting vertical transmission. It is known that the virus is capable of inducing several cytogenetic aberrations in peripheral blood (STOCCO DOS SANTOS et al., 1998), with molecular diagnosis being an essential tool to guarantee that there will be no clastogenicity induction by the virus and interference in other tests.

### 3.4. Comet (EC) assay in blood cells

The results of the comet assay are shown in table 5.

**Table 5.**
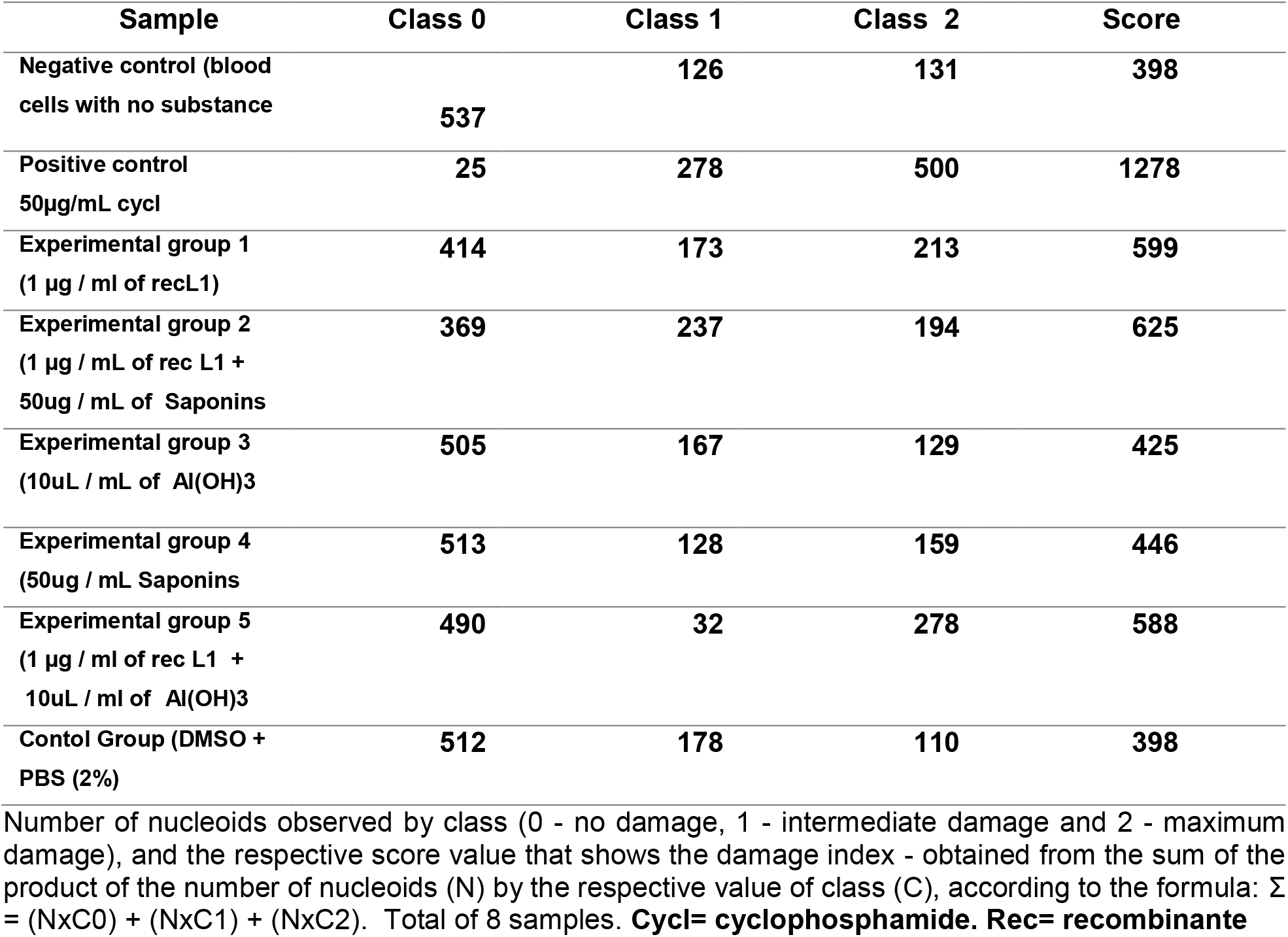
Results and scores obtained in the analysis of mutagenic potential by comet assay from 8 sample.

Based on the score values shown in table 5, the Kruskal-Wallis test was performed with a significance level of 5%. The test showed statistically significant differences (p = 0.0004) between the groups. Based on this result, Dunn’s post-hoc test was performed, also with a 5% significance level shown in figure 8, which revealed significant statistical differences between the negative and positive control group, as well as between positive control and the group treated only with Aluminum Hydroxide, Saponins and the DMSO diluent, however, the test did not reveal significant differences between the positive control group and the one treated with the combination of L1 protein and adjuvants, nor the group treated with L1 protein alone. In these cases, there were also no significant differences with the negative control, which represents intermediate values as shown in table 6.

**Table 6.** Dunn’s post-hoc test statistical values obtained using the GraphPad Prism software version 5

| Dunn's Multiple Comparison Test | Difference in rank sum | Significant? P < 0.05? |
| --- | --- | --- |
| C+ vs C- | 37.81 | Yes |
| C+ vs S | 34.25 | Yes |
| C+ vs L1+S | 22.50 | No |
| C+ vs L1 | 26.38 | No |
| C+ vs DMSO | 38.25 | Yes |
| C+ vs H | 36.25 | Yes |
| C+ vs L1H | 23.56 | No |
| C- vs S | -3.563 | No |
| C- vs L1+S | -15.31 | No |
| C- vs L1 | -11.44 | No |
| C- vs DMSO | 0.4375 | No |
| C- vs H | -1.563 | No |
| C- vs L1H | -14.25 | No |

The comparison of the score values between the groups is shown in figure 2.

**Figure 2.**
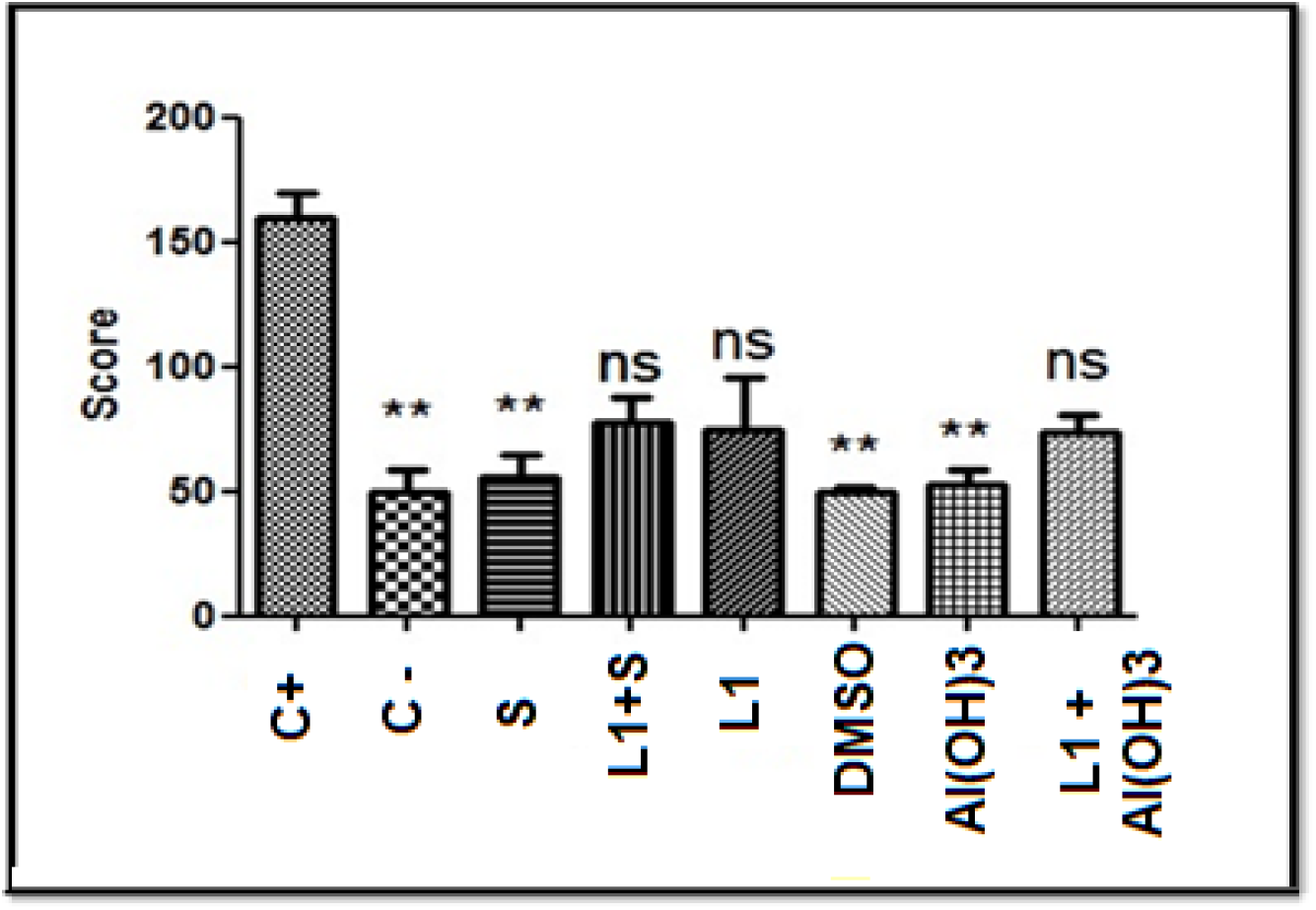
Statistical graph comparing the score values of the treated samples in relation to the positive control, in which the C-, Saponins, DMSO and H groups showed significant differences with the comparative group, not showing a mutagenic character.

### 3.5. Micronucleus Test with Cytokinesis Block (TMBC) in Peripheral Blood

The test was done repeatedly and was not satisfactory, as there were not enough cells to count, since more than 80% were hemolyzed, including in the negative control, which leads us to believe that it is not something related to the toxicity of the drugs applied . Figure 3 shows a comparison between cells without the presence of micronuclei and micronucleated cells.

**Figure 3.**
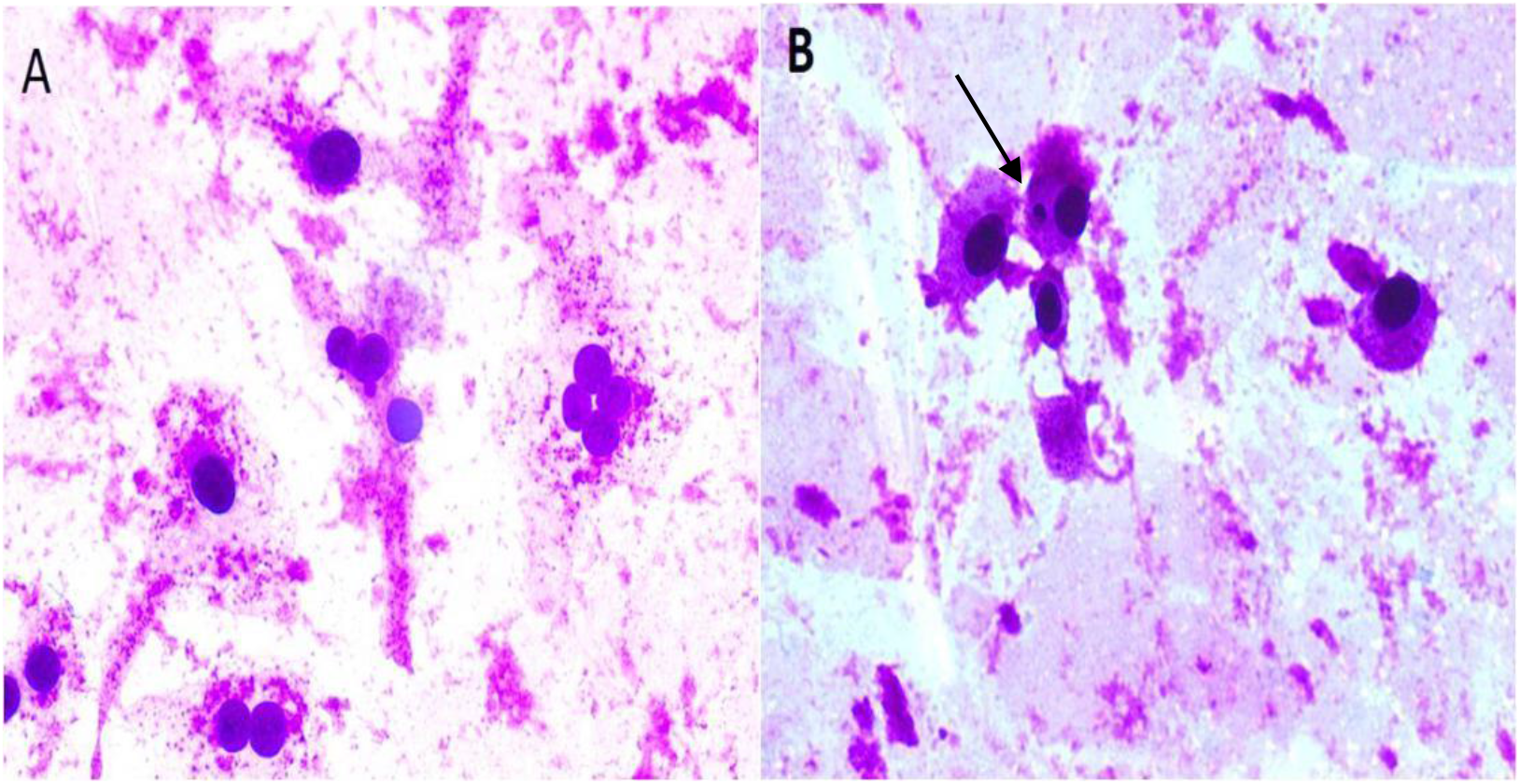
Microscopic analysis of the Micronucleus Test in blood cells **A)** Lymphocytes without the presence of a micronucleus; **B)** Lymphocyte with micronucleus.

### 3.6. Comet (EC) assay in cell culture

The results observed in the analysis of the mutagenic potential by comet assay in Vero cells are shown in table 7.

**Table 7.** Number of nucleoids observed by class (0 - no damage, 1 - intermediate damage and 2 - maximum damage), and the respective score value that shows the damage index - obtained from the sum of the product of the number of nucleoids (N) by the respective value of class (C), according to the formula: S = (NxC0) + (NxC1) + (NxC2).

| Sample | Class 0 | Class 1 | Class 2 | Score |
| --- | --- | --- | --- | --- |
| Negative control<br>Only Vero cells | 140 | 05 | 03 | 13 |
| Positive control (50 µg /<br>mL cycl | 17 | 40 | 93 | 226 |
| Experimental group 1<br>1 µg / mL of rec L1 | 61 | 37 | 53 | 141 |
| Experimental group 2<br>(1 µg / ml of rec L1 +<br>50ug / ml of Saponins) | 63 | 29 | 58 | 145 |
| Experimental group 3<br>(10uL / mL of Al(OH)3 | 61 | 33 | 53 | 138 |
| Experimental group 4<br>(50ug / mL Saponins) | 33 | 54 | 63 | 180 |
| Experimental group 5<br>(1 µg / ml of rec L1 +<br>10uL / ml of Al(OH)3 | 60 | 41 | 59 | 159 |
| Control Group DMSO +<br>PBS (2%) | 112 | 21 | 19 | 59 |

Based on the score values shown in table 7, the Kruskal-Wallis test was performed with a 5% significance level as shown in table 8. The test showed statistically significant differences ( $p = 0.0038$ ) between the groups. Based on this result, Dunn's post-hoc test was performed, also with a 5% significance level. **Cycl= cyclophosphamide. Rec= recombinante**

The test revealed significant statistical differences between the negative and positive control group, as well as between this and the group treated only with the DMSO diluent, however, the test did not reveal significant differences between the positive control group and the one treated with the L1 protein combination and the adjuvants that were the interest groups, nor with the separate drugs, but there was also no significance with the negative control, being an intermediate result, according to table 8.

**Table 8.** Dunn’s post-hoc test statistical values obtained using the GraphPad Prism software version 5

| Dunn's Multiple Comparison Test | Difference in rank sum | Significant? P < 0.05? |
| --- | --- | --- |
| C+ vs C- | 21.00 | Yes |
| C+ vs S | 3.667 | No |
| C+ vs L1+S | 9.833 | No |
| C+ vs H | 12.33 | No |
| C+ vs L1 | 12.67 | No |
| C+ vs LI+H | 6.500 | No |
| C+ vs DMSO | 18.00 | Yes |
| C- vs S | -17.33 | No |
| C- vs L1+S | -11.17 | No |
| C- vs H | -8.667 | No |
| C- vs L1 | -8.333 | No |
| C- vs LI+H | -14.50 | No |
| C- vs DMSO | -3.000 | No |

The comparison of the score values between the groups is shown in figure 4.

**Figure 4.**
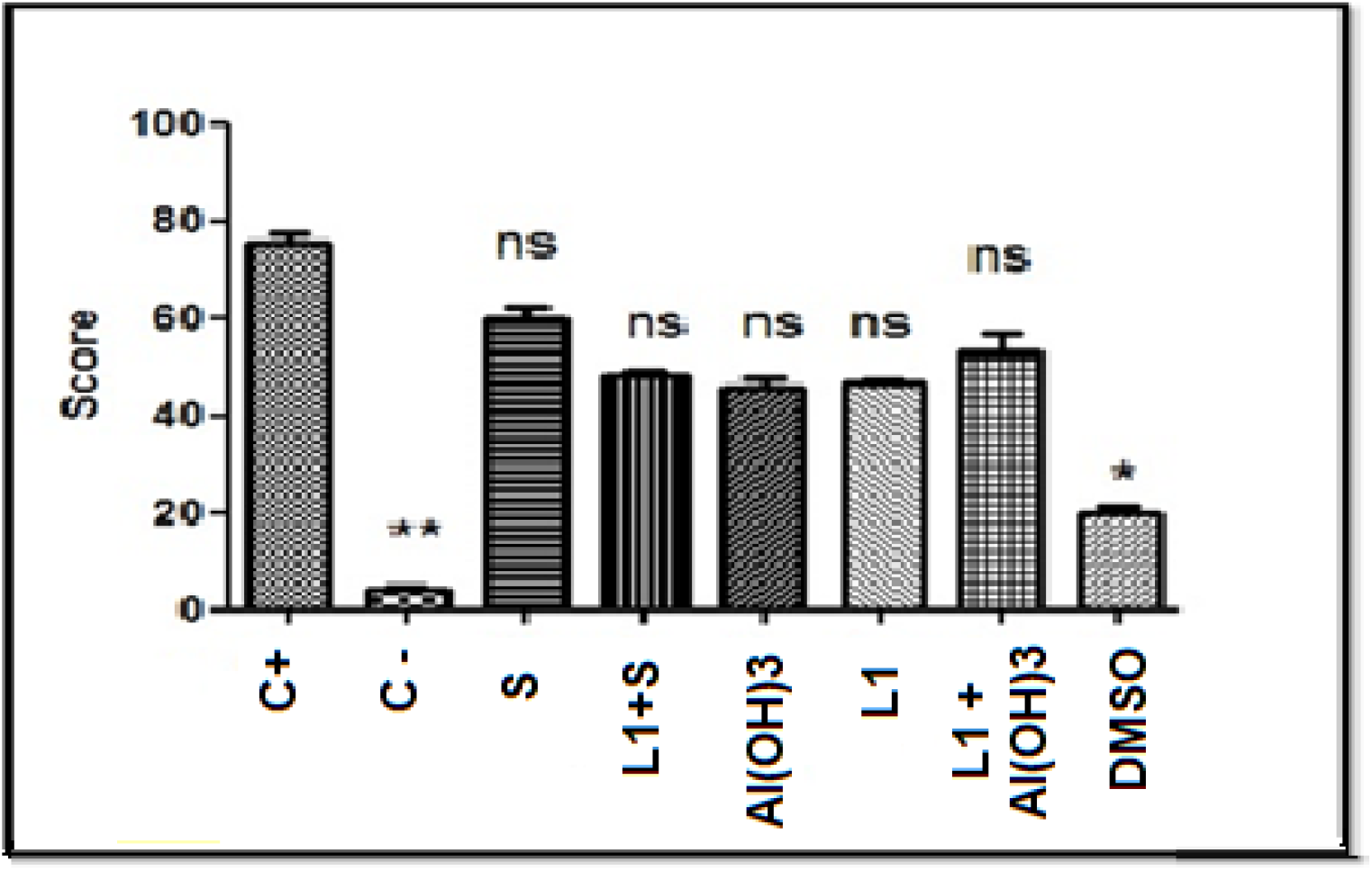
- Statistical graph comparing the score values of the treated samples in relation to the positive control, in which only the C- and DMSO groups showed significant differences with the comparative group.

### 3.7. Micronucleus Test with Cytokinesis Block in Cell Culture

1,000 cells per slide were analyzed. Analyzes of TMN in Vero cells were performed by evaluating the frequency of micronucleus formation (MNr0 = a / b, where: a - number of cells with MN and b - total number of cells).

The values found in the micronucleus test are shown in table 9.

**Table 9.** Number of micronuclei found per slide and calculated frequency

| Samples | Nº Micronuclei | MN<sub>r0</sub> |
| --- | --- | --- |
| **C+** | 691 | 0, 691 |
| **C-** | 77 | 0, 077 |
| **Al(OH)<sub>3</sub>** | 148 | 0, 148 |
| **L1+ Al(OH)<sub>3</sub>** | 176 | 0, 176 |
| **L1** | 106 | 0, 106 |
| **Saponin** | 226 | 0, 226 |
| **L1+Saponin** | 266 | 0, 266 |
| **DMSO** | 113 | 0, 113 |

**Table 10.**
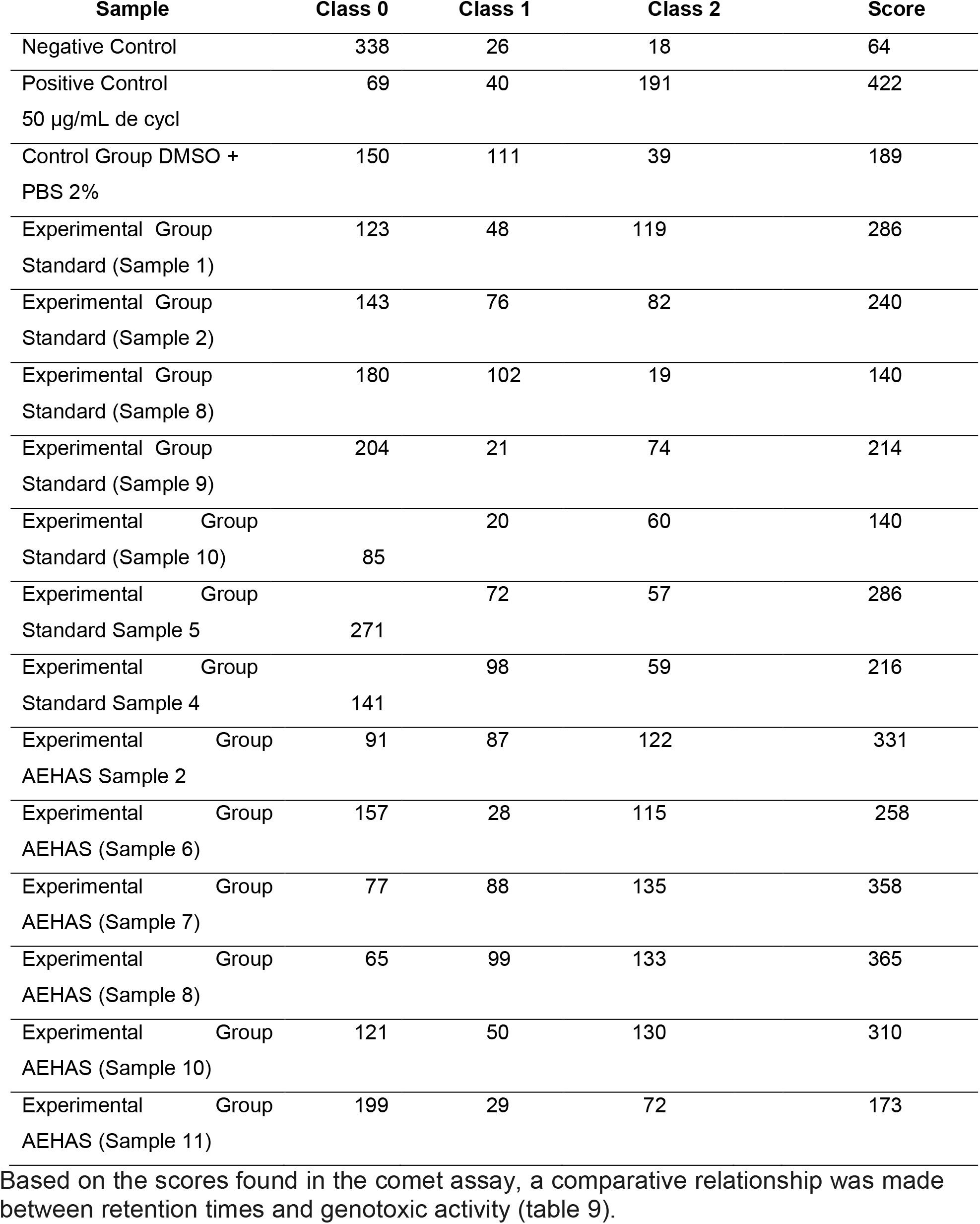
Number of nucleoids observed by class (0 - no damage, 1 - intermediate damage and 2 - maximum damage), and the respective score value that shows the damage index - obtained from the sum of the product of the number of nucleoids (N) by the respective value of class (C), according to the formula: S = (NxC0) + (NxC1) + (NxC2). From 3 samples Based on the scores found in the comet assay, a comparative relationship was made between retention times and genotoxic activity (table 9).

In Figure 5 is showed a comparative histogram of the micronucleus percentages counted among the samples with the 8 different treatments in Vero cells.

**Figure 5.**
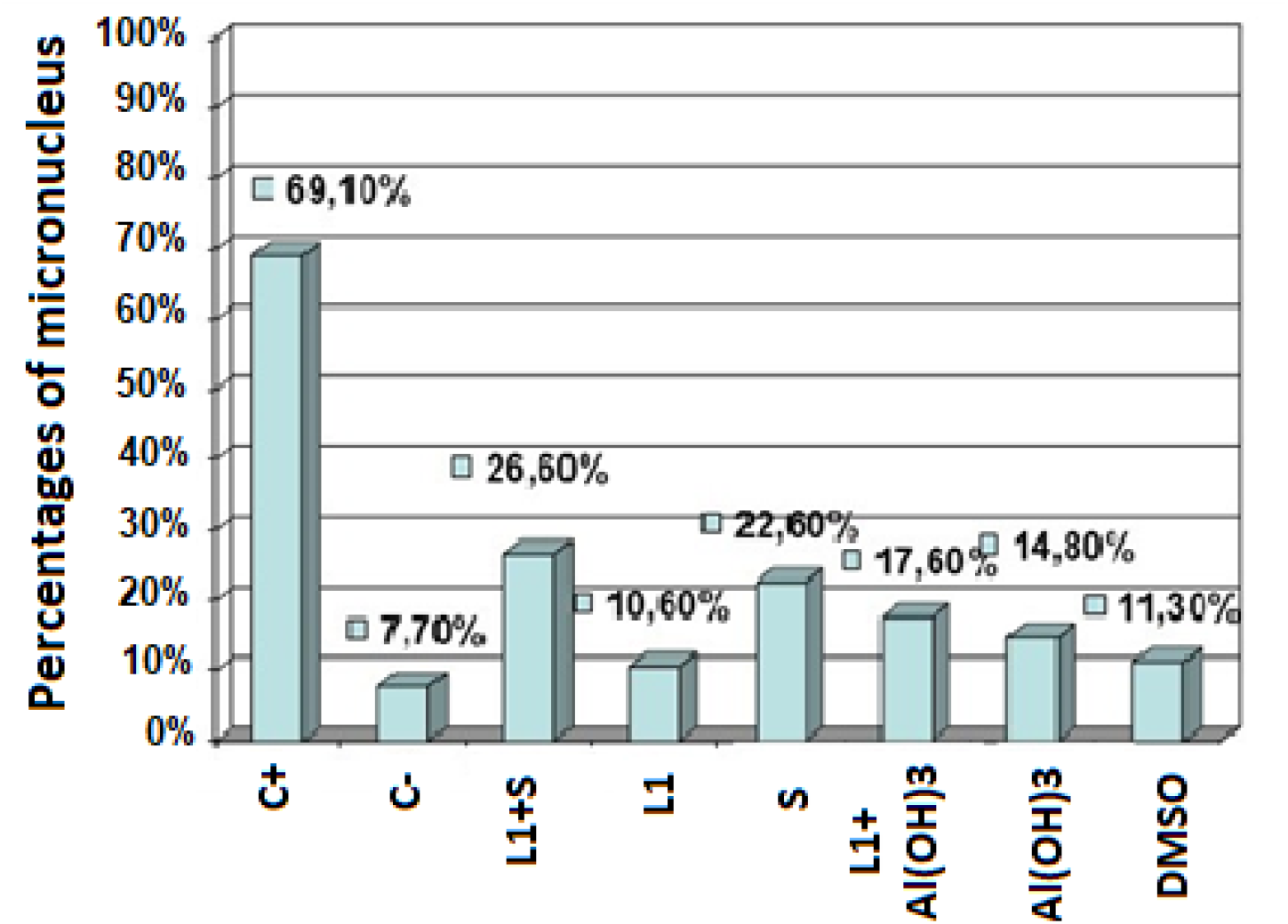
Comparative histogram of the percentages of micronuclei counted among the samples with the 8 different treatments in Vero cells.

Based on these values, it can be considered that 7.7 of micronuclei found in the negative control of a total of 1,000 cells analyzed are part of the basal cell metabolism and therefore this value was subtracted from the other percentages, being found for the combination of L1 with saponins an increase of 18.9% of micronuclei to the negative control and 9.9% when applying the combination of L1 with Aluminum Hydroxide, almost double the addition of micronuclei.

The values approximate the sum of the percentages of separate drug micronuclei, with 2.9% for L1, 14.9% for saponins, 7.1% for Aluminum Hydroxide and 3.6% for DMSO. No value came close to the positive control.

### 3.8 Flow cytometry: assay with Annexin V-PI

The data were analyzed using the BD Accuri C6 software. Statistical analysis was performed based on the average percentage of living cells, using the two-way ANOVA test followed by the Tukey post-hoc test (P <0.05) using the GraphPad Prism software version 5 (GraphPad Software, Inc.) according to figure 6.

**Figure 6.**
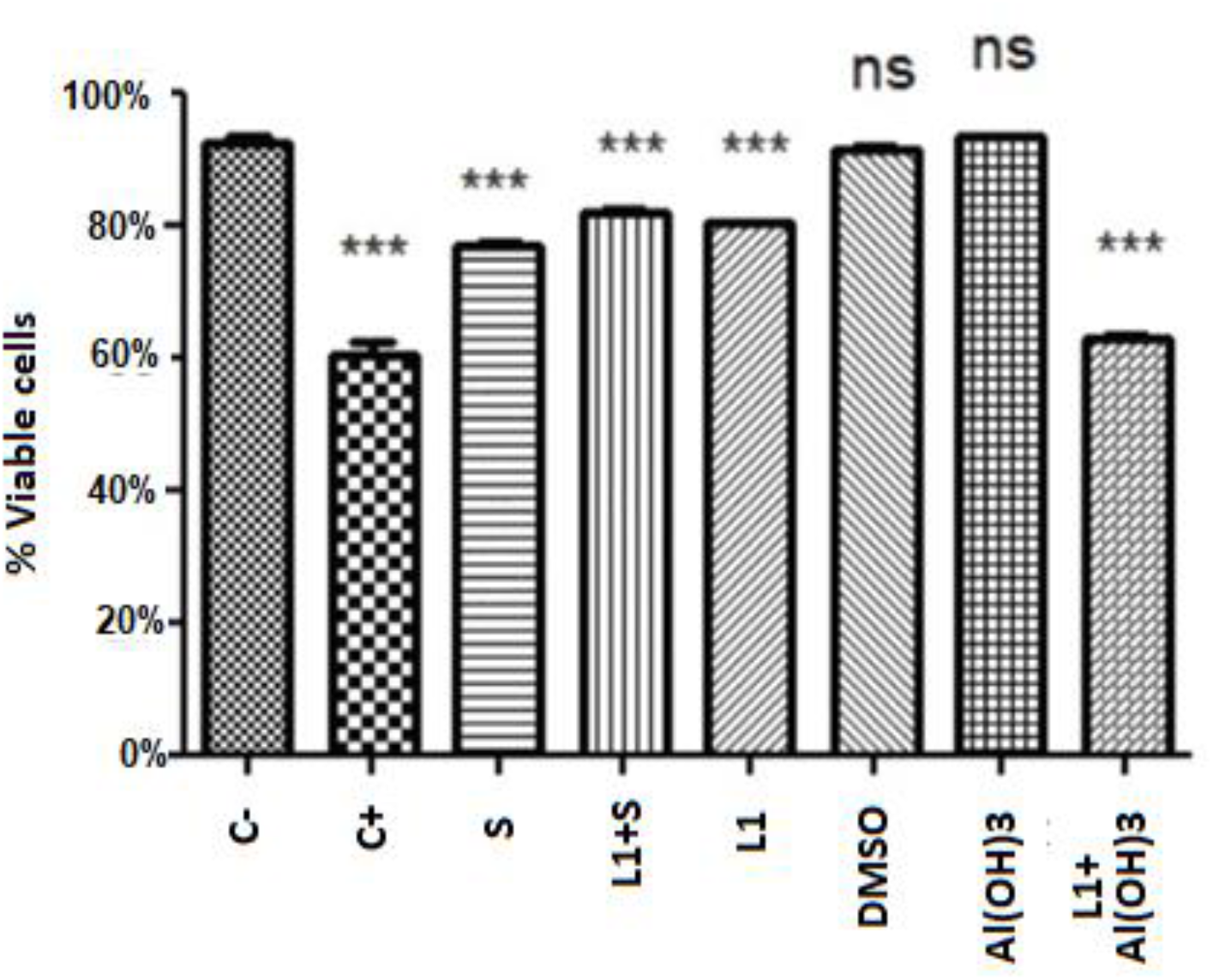
Statistical graph comparing the means with standard deviation of viable cells of the treated samples in relation to the negative control, in which only the H and L1 + H groups did not show significant differences with the comparative group.

Statistical analysis was performed based on the average percentage of living cells. The two-way ANOVA test followed by the Tukey post-hoc test showed statistically significant differences (p <0.0001) between the groups, also with a 5% significance level.

All mean values showed high significant differences in relation to viable cells with the negative control, except for the conjugate of L1 protein with Aluminum Hydroxide and Aluminum Hydroxide when applied alone, which presented values higher than the negative control.

Figure 7 shows a dot plot graph of the intersection between the annexin fluorescence axes V-FITC and PI where the viable cell population is represented in the lower left quadrant (Annexin V-PI-), the early apoptotic cells in the lower right quadrant (Annexin V + PI-), late apoptotic / necrotic cells in the upper right quadrant (Annexin V + PI +), dead cells (Anexin V - PI +) in the upper left quadrant.

**Figure 7.**
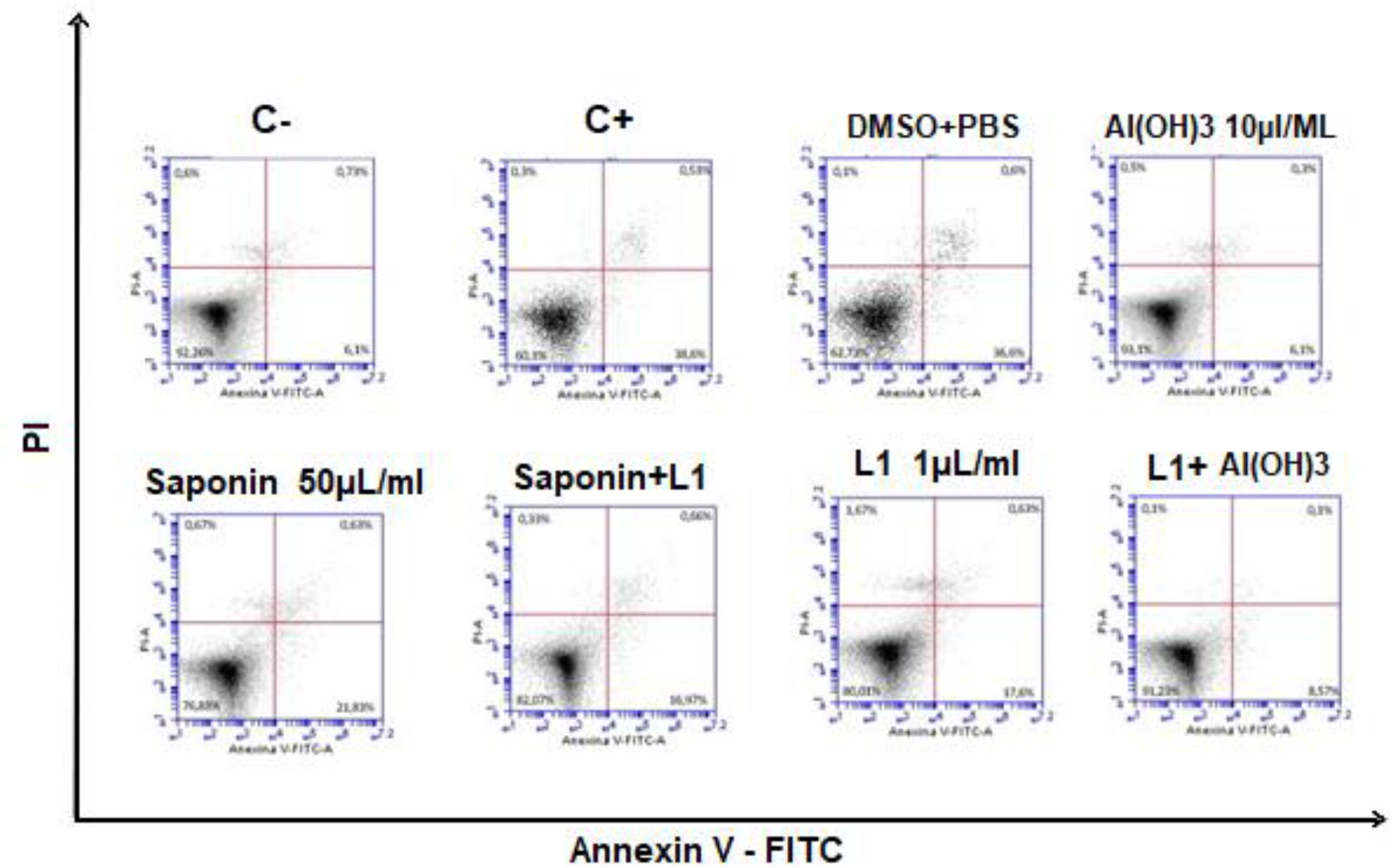
Dot plot graph of the data analyzed using the BD Accuri C6 software for Vero cells treated according to Table 3 based on the average percentage of live cells, using the two-way ANOVA test followed by the test Tukey post-hoc (P <0.05) using GraphPad Prism software version 5 (GraphPad Software, Inc.)

In figure 8, the data obtained from the triplicates are represented in a statistical graph based on the average percentage of cells in each phase and standard deviation, using the two-way ANOVA test followed by the post-hoc Bonferroni test (p <0.05) using the Graphpad prism software version 5 (Grandpad software, inc.).

**Figure 8.**
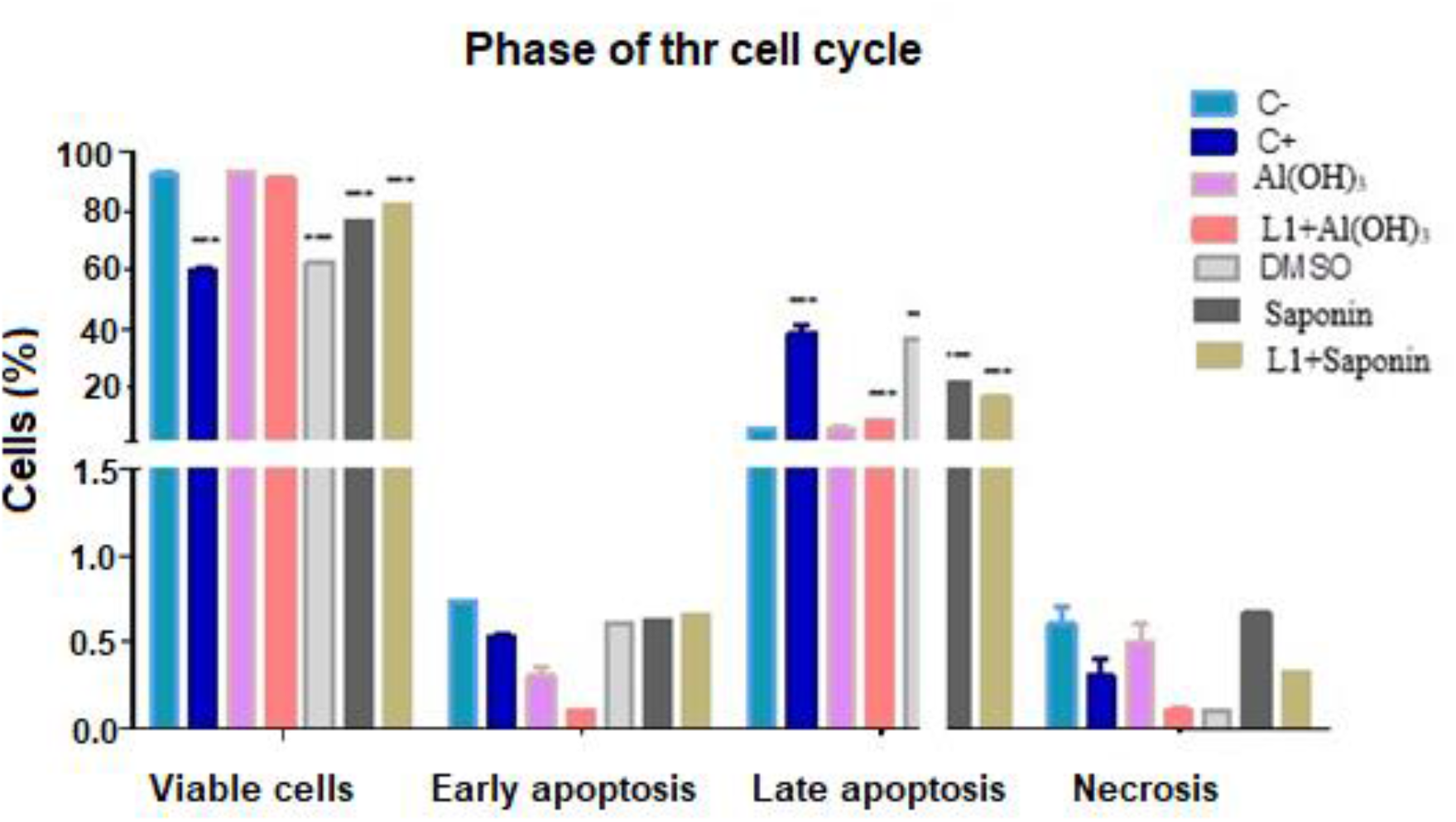
Statistical graph of the data analyzed using the BD Accuri C6 software for vero cells treated according to table 3 in comparison to cells without treatment (C-). Based on the average percentage of cells distributed in each phase of the cell cycle and standard deviation, the two-way ANOVA test was performed followed by the Bonferroni post-hoc test (p <0.05) using the Graphpad prism software version 5 (GrandPad Software, Inc.).

### 3.9. Isolation of substances present in the Sigma standard saponin extract and in the Saponin extract under test by chromatographic fractionation

At Figure 9 is showed the results obtained in the isolation of the substances present in the extract of saponin Standard Sigma and in the extract Saponin test, fractionated and identified by HPLC. The eluted fractions and with similar retention times, had their genotoxicities evaluated for later characterization by mass spectrometry. This step resulted in the separation of 21 fractions. Of these 21 fractions, 13 were chosen, 7 from the standard and 6 from the extract (indicated by the arrow in figure 9).

**Figure 9.**
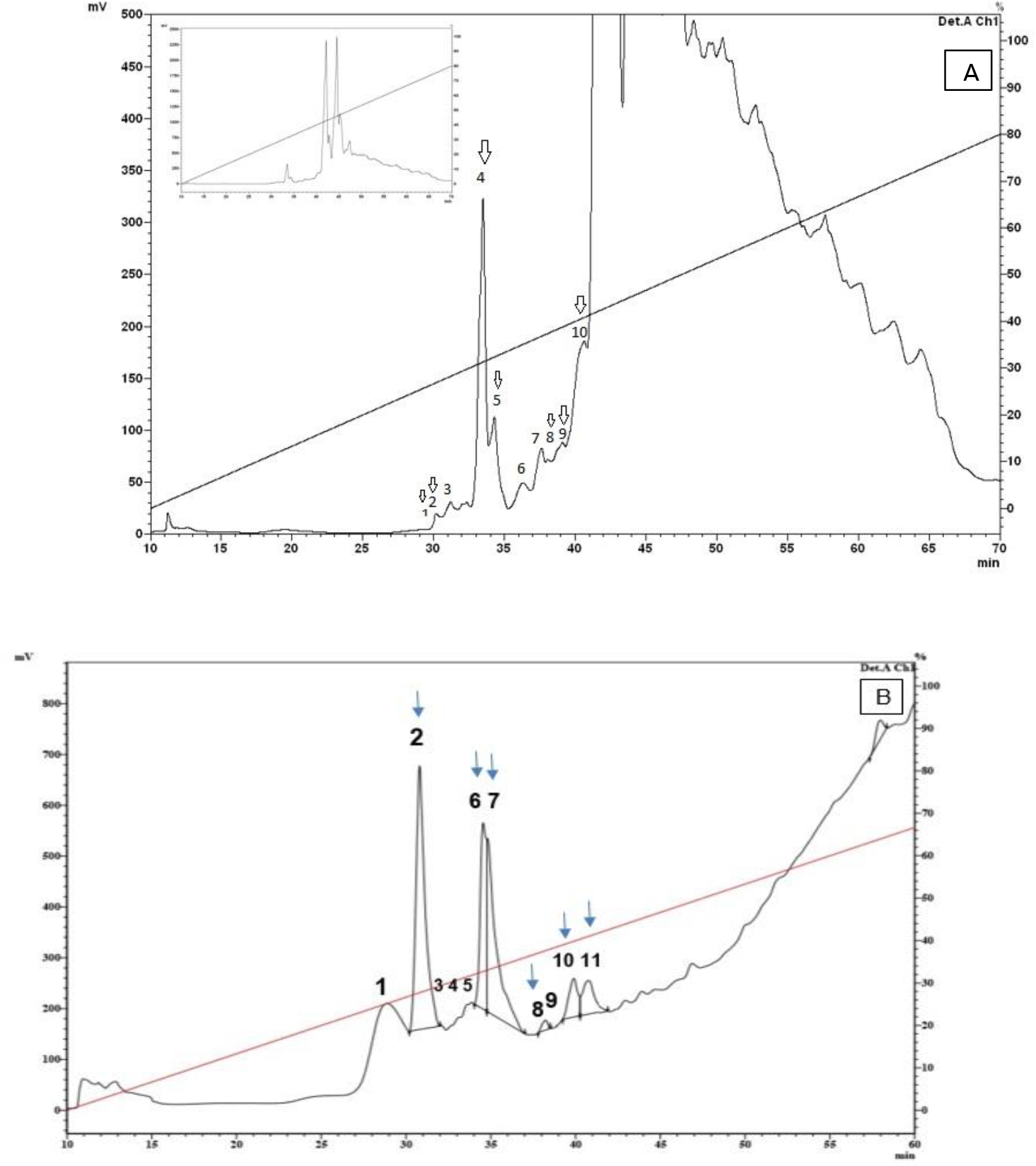
Chromatographic analysis of the sigma Standard saponin extract (A) and the Saponin extract under test (B), in a UFLC Shimatzu system, model Proeminence. 21 fractions were eluted, which were collected and selected 13 (indicated by arrow). The material was fractioned in a Jupiter C18 semi-preparative reverse phase column (250 mm x 10 mm, Phenomenex) with a linear gradient from 0 to 80% acetonitrile in acidified water for 60 minutes, in a flow of 2mL / min. Absorbance was monitored at 225 and 280 nm.

### 3.10. Comet assay (EC) in cell culture of the fractions obtained

In table is showed the number of nucleoids observed by class in test comet performe with differents sample in Vero cells

In Table 11 is showed the comparison between standard and sample in relation to retention time and genotoxic activity scores of the comet assay.

**Table 11.** Comparison between standard and sample in relation to retention time and genotoxic activity scores of the comet assay.

|  |  |
| --- | --- |
| Negative Control | Score: 64 |
| Positive Control | Score: 422 |
| DMSO | Score: 189 |

| Standard nº Pico | Retencion time (min) | score |
| --- | --- | --- |
| 1 | 28,13-30,54 | 286 |
| 2 | 30,54-31,66 | 240 |
| 4 | 32,64-34,13 | 216 |
| 5 | 34,13-35,22 | 286 |
| 8 | 38,28-39,28 | 140 |
| 9 | 39,28-40,84 | 214 |
| 10 | 40,84-42,57 | 140 |

| EHAS nº Pico | Retencion time (min) | Score |
| --- | --- | --- |
| 2 | 30,17-32,29 | 331 |
| 6 | 34,90-35,11 | 258 |
| 7 | 35,11-37,14 | 358 |
| 8 | 37,14-38,58 | 365 |
| 10 | 39,47-41,42 | 310 |
| 11 | 41,42-42,04 | 173 |

Four fractions (two from the standard and two from the extract sample) that demonstrated similarity in relation to the retention time in the chromatography and genotoxic activity in the comet assay were selected for structural analyzes in mass spectrophotometry, so that there was a confirmation that they were the same substance and enable a first study of separation of the compound that would be causing genotoxic effect and impairing the use of saponin as an adjuvant in this respect.

### 3.11 Analysis and characterization of substances by mass spectrometry

The data obtained in mass spectrometry were analyzed separately using the Xcalibur 2.0 software (Thermo Electron, USA). The Mass Analyzer 1.03 software was used to confirm the average masses of the fragments and the deconvolution of the spectra loads.

The fractions of extract nº 06 and nº 10 resulted in 740,14Da and 740,15Da respectively, and the fractions of standard nº5 and nº9 resulted and mass values of 740,09Da and 740,17Da (figures 10 and 11), being probably the same substances. It was not possible to identify the substances exactly due to the lack of a database available for this class of compound.

**Figure 10.**
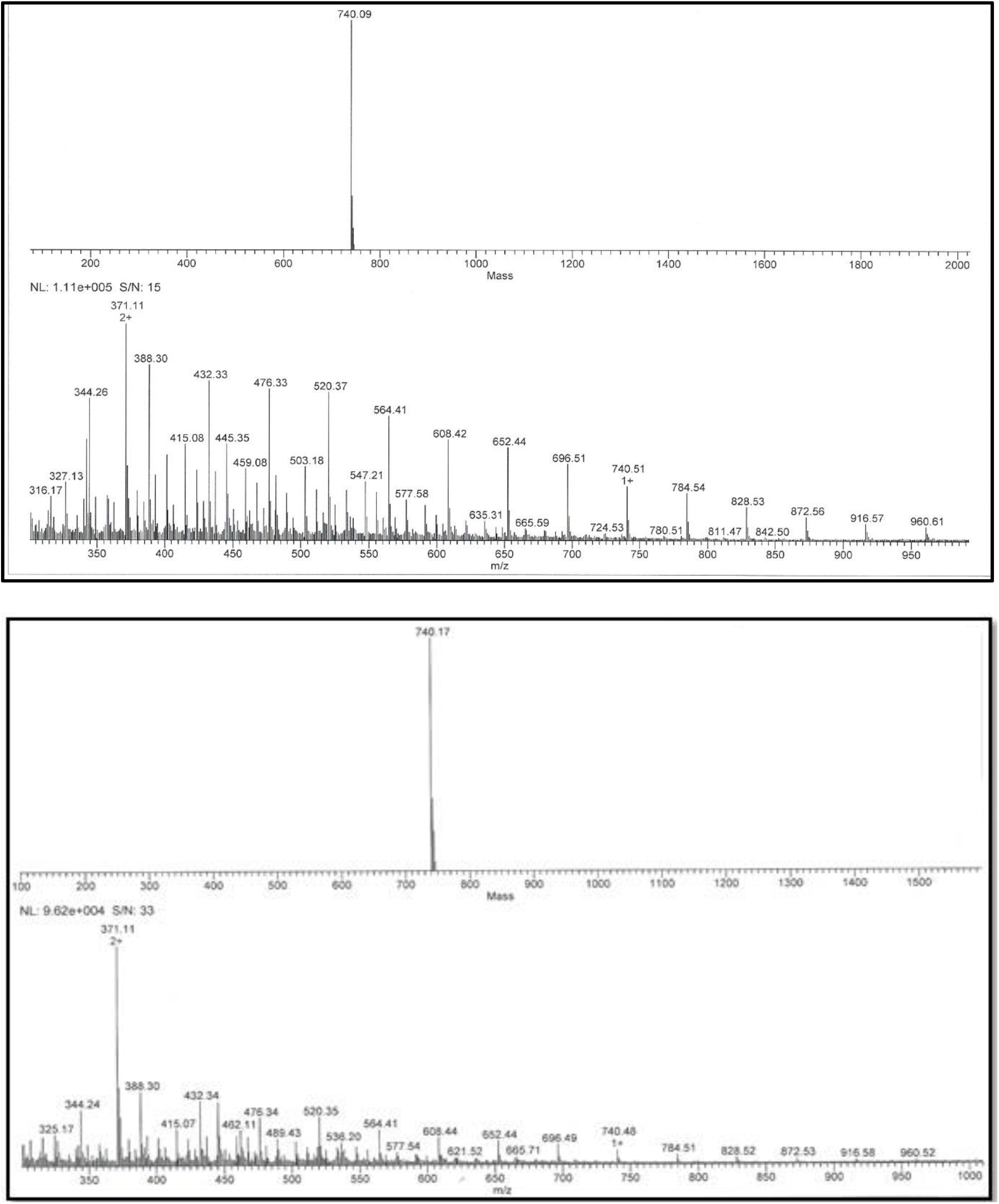
Deconvolution of the spectra of the fractions of pattern nº05 and nº09. The images were obtained through analysis using the Mass Analyzer 1.03 software. Through the deconvolution of the ions (m / z), the molecular masses of 740.09 Da for fraction nº05 and 740.17 Da for fraction nº09 of the standard

**Figure 11.**
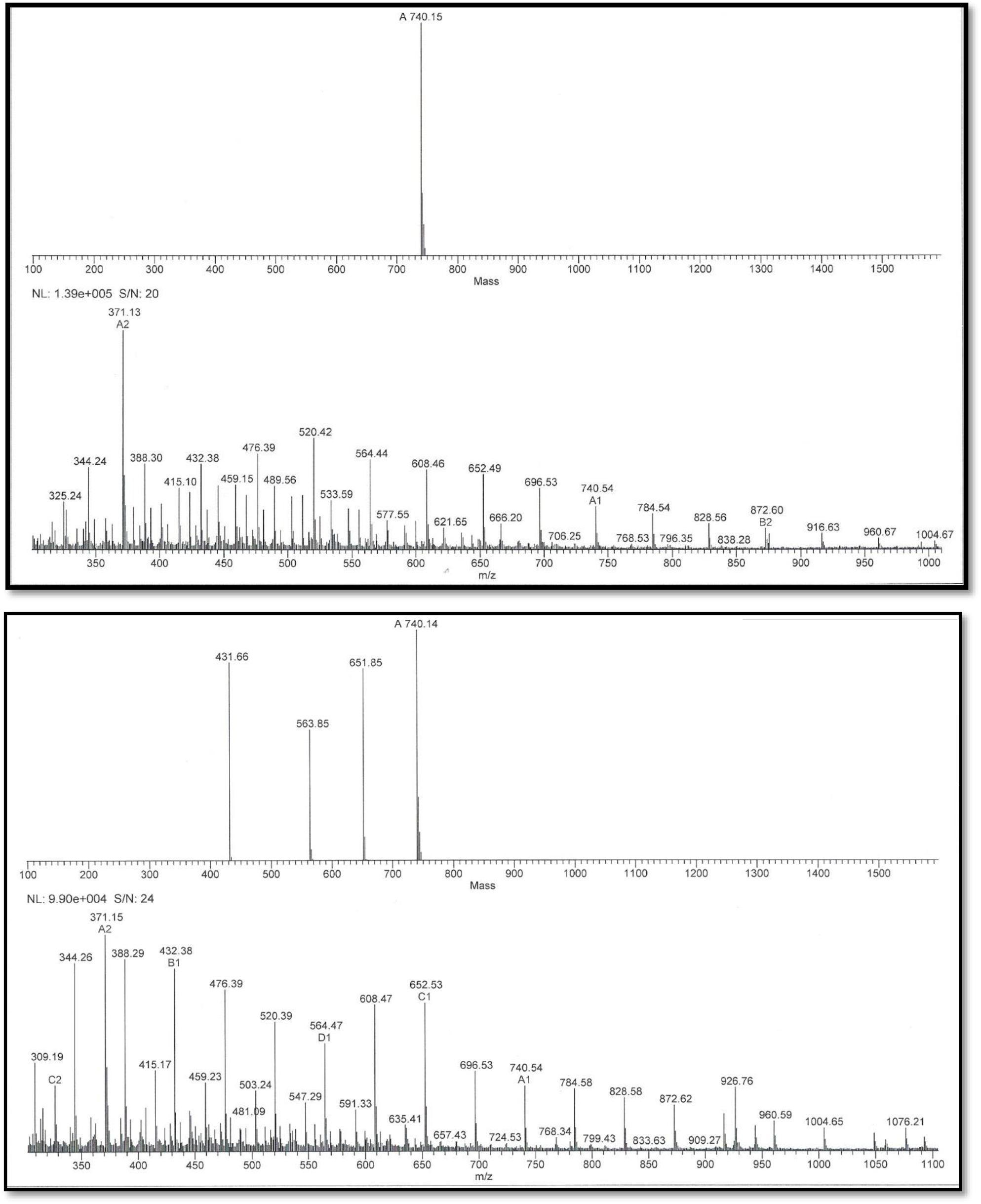
Deconvolution of the spectra of the fractions of sample nº06 and nº 10. The images obtained through analysis by Mass Analyzer 1.03 software. Through the deconvolution of the ions (m / z), the molecular masses of 740.15 Da for fraction No. 06 and 740.14 Da for fraction No. 10 of the standard

## 4. DISCUSSION

Vaccines are great allies in combating the actions of infectious agents. Its effectiveness has been verified in both prophylactic and therapeutic aspects. To increase this effectiveness, several substances have been administred together with the vaccine product. These substances are known as adjuvants, from the Latin adjuvare (to improve) (SCHIJNS, 2000). Different adjuvants have been used: mineral (aluminum hydroxide - Alum), vegetable (saponins), emulsions (incomplete Freund’s adjuvant) and even derivatives of infectious agents (Freund’s complete adjuvant). (RAMON, 1925), however, the local and / or systemic toxicity conferred by these adjuvants has been a great challenge to license a new component (SUN et al., 2009).These substances provide greater immunological action for vaccine products, which is of utmost interest in terms of public health and biotechnology. However, although adjuvants have been used for decades and there are reports of adverse reactions, there are still no studies on possible cytotoxic actions of these products (GUPTA & SIBER, 1995). Bovine papillomavirus (BPV) is the etiological agent of bovine papillomatosis, an infectious disease characterized by the presence of papillomas that regress spontaneously, but that can persist and progress to malignancy, affecting meat and milk production segments, which represents a serious problem of livestock considering that at least 60% of cattle are contaminated in Brazil (STOCCO DOS SANTOS et al., 1998). Although management control is seen as a measure of disease control, there is a need to develop new vaccine and diagnostic technologies, since the virus can be transmitted vertically (ROPERTO et al., 2012).

However, like all vaccines based on recombinant proteins, the final formulation requires the use of substances capable of improving the immune response against a specific antigen (REED et al., 2009). In addition, by increasing the immune response, adjuvants reduce the necessary amount of antigen in the final formulation of the vaccine, reducing its cost (AGUILAR et al., 2007).

Mineral compounds (calcium phosphate, aluminum hydroxide) are widely used in human and veterinary vaccines. The use of aluminum hydroxide, also known as Alum, was introduced in 1926 (GLENNY et al., 1926). However, Alumem is a poor inducer of T cell response, an important characteristic in the development of a vaccine. In this scenario, saponins emerge as a promising candidate for adjuvant for veterinary use (PETROVSKY et al, 2004), due to their greater adjuvant potential in relation to alum (GUPTA et al., 1995).

In view of the antigenicity of L1 protein and the remarkable adjuvant capacity of saponins, the present study aimed to evaluate the mutagenic and genotoxic potential of isolated and purified protein, as well as its effects when associated with saponins and also to compare it to hydroxide aluminum, using in vitro tests.

The levels of clastogenicity were detected by the comet assay and the genotoxic potential, through the micronucleus test. The degree of apoptosis was detected by flow cytometry using Annexin-PI.

Together, these methods allow to evaluate the mutagenic and cytogenetic potential of the formulations tested, which is a need required by regulatory agencies, since toxicity limits the release and use of drugs and these studies serve as indicators for conducting pre-clinical studies, and should be performed in cells and later in animals (MARTIN et al., 2009).

Such tests were carried out on Vero cells and bovine peripheral blood, since According to the International Workshop on Genotoxicity Test Procedures (TICE et al., 2000), at least two independent cell cultures must be tested successively, in addition to Vero cells have been widely used, not only for microbiology, but also for the production of vaccines for human use (OSADA et al., 2014), in addition to which this strain does not have oncogenic properties, as has been demonstrated in several studies, being accepted by the Organization World Health Organization - WHO - for use as a substrate including in the production of vaccines for humans (FURESZ et al., 1989; VINCENT FALQUET et al., 1988; HORAUD, 1992).

The results obtained by the comet assay, a sensitive method for studying DNA damage in individual cells (SINGH et al., 1988; SARAN et al., 2008), in peripheral blood, showed that the negative control group had a lower number of comets. This result was expected, since the biological material was transported and processed after three hours of its collection, reducing the interference of exogenous environmental factors, in addition, the blood samples, as verified in the PCR assay, were not infected by BPV, reducing thus the influence of endogenous environmental factors that could influence the results.

The positive control group had a high number of class 2 comets, a result that was also expected, as it was treated with 50 ug / mL of cyclophosphamide, chemotherapy and mutagenic, cytotoxic and teratogenic immunosuppressants (BRITO et al., 2008).

Dunn’s post-hoc test did not reveal statistical differences between some groups tested compared to the positive control, pointing out cytogenetic damage compatible with the samples treated with the L1 protein, the vaccine compositions composed of L1 conjugated saponins and L1 with Aluminum Hydroxide, the which suggested that L1 in these cases is the determining factor of cytogenicity.

These groups also did not present significant differences with the negative control, acquiring an intermediate character between the two controls.

DMSO, aluminum hydroxide, saponins and negative control showed to be significantly different from the positive control and did not show significant differences with the negative control.

In the data obtained in Vero cells, the results were similar to those of peripheral blood for these treatments. However these cells were more sensitive to treatment with saponins and aluminum hydroxide alone, which gave them cytogenicity, being only the diluent DMSO and the negative control significantly different from the positive control.

No group showed a significant difference with the negative control, and the treated samples also acquired an intermediate character between the two controls.

The micronucleus test has been widely used as a primary test in toxicological genetics for its ability to assess genotoxicity in a simple and fast way, detecting chromosomal damage induced as a mutagenicity biomarker (HEDLE, et al. 1983).

In tests with L1 protein an increase of 2.9% of micronuclei was observed, in the combination of L1 with saponins there was an increase of 18.9% of micronuclei compared to the negative control, and 9.9% when applying the combination of L1 with Aluminum Hydroxide, which represents almost twice the number of micronuclei, but no sample was close to the value of the positive control, which added 61.4% of micronuclei in relation to the negative control which represented only 7.7% of micronuclei a total of 1000 cells counted. The diluent DMSO added 3, 6%, Aluminum hydroxide 7, 1% and Saponin only, 14.9% of micronuclei.

The sum of the percentages of micronuclei found in the application of separate drugs (Aluminum Hydroxide or Saponins) with the percentage of micronuclei found in the application of L1, also separated, approximates the percentage found of micronuclei in L1 vaccine formulations with Aluminum Hydroxide and L1 with Saponins, the latter being at a disadvantage slightly increasing the formulation’s genotoxic potential. No value came close to the positive control.

The DNA repair system is essential for maintaining genetic stability and consequently for maintaining life and the importance of these repairs is evident, given the huge investment made by cells in enzymes for this purpose (FRIEDBERG et al., 2005). When DNA damage occurs, checkpoint mechanisms are activated, triggering a cellular response that triggers a break in the cell cycle. Then, depending on the type, degree of damage and the physiological state of the cell, the cell response can happen in two ways: by activating the DNA repair system, resulting in the restoration of its original chemical structure, or, if the damage the DNA are very severe, and therefore not liable to repair, the cell chooses to eliminate itself through apoptosis (MASLOV and VIJG, 2009).

In this context, the annexin V-PI assay, the annexin V-PI assay pointed out that none of the treated samples had apoptotic potential similar to the positive control, since there was a statistically significant difference, with the exception of DMSO, which showed little statistical difference, but still significant.

However, in relation to the negative control, samples with Saponins, Saponins with L1, DMSO and L1 showed significant differences, being in an intermediate state of cell viability between positive and negative control. The vaccine combination of Aluminum Hydroxide with L1 and only Aluminum Hydroxide did not show significant differences with the negative control, and, analyzing the values of viable cell means, aluminum hydroxide increased the number of viable cells in relation to the negative control when applied; there is no report of this protective capacity of aluminum in the literature.

The apoptotic capacity of saponins can be attributed to the presence of surfactant, which binds to the cholesterol present in the cell membrane, leading to the formation of pores and hemolysis (O’HAGAN, 2003). Among biological activities, hemolysis is the characteristic common to many saponins with different structures. This activity is due to the fact that saponins in general form complexes with the steroids of the erythrocyte membrane, causing an increase in permeability and the consequent loss of hemoglobin (COSTA, 2001; WANG et al., 2007).

The hemolytic effect of saponins is greatly influenced by the polar nature of substituents linked to aglycon (VOUTQUENNE et al., 2002), as well as their immunoadjuvant activity (SOLTYSIK et al., 1995), however the evidence is not yet conclusive, besides most studies focus on the immunoactivity of the Quillaja saponin (KENSIL et al., 1995), which is only one among many existing and, therefore, differs in the chemical structure, requiring further studies with other species.

The results also suggest toxicity related to the DMSO diluent, which showed no significant difference with the positive control. DMSO was chosen as a saponin diluent, due to its interesting pharmacotechnical and pharmacological properties for a vaccine, as it has an intense penetration capacity, facilitating the carrying of many substances associated with it through membranes (BLYTHE et al., 1986; RAND-LUBY et al., 1996) and is myorelaxative, associated with tranquilizing and sedative effects, an effect that has been observed in several species (ROSENBAUM, 1965; BRAYTON, 1986;). This property undoubtedly results mainly from the comfort resulting from other properties, that is, anti-inflammatory and analgesic (BLYTHE, 1986; BRAYTON, 1986).

In a recent review (2020), O’HAGAN et al has showed that despite of recombinant antigens be very safe, its potency, in general, is lower that convencional vaccins. So, to improved their potency, the interest in news adjuvants has increased. However, the tolerability profile of new candidate adjuvants is very different to tradicional adjuvante as Alum, including many practical limitations with these new adjuvants as ” difficulties in the reliable supply of materials of appropriate quality and consistency, at suitable quantity, formulation robustness and process reproducibility”. Thus, the development of new adjuvants can be a limitant factor to obtaining of some new vacines. Because of this, for the development of new generation vaccines, an extensive study of the traditional adjuvants already available, as well as the development of new and safe adjuvants are essential

Novertheless, adjuvants can play an important role in developing of a new vaccines as COVID-19 vaccine. Recently a Novavax’s COVID-19 Vaccine Clinical Trials was performed using as adjuvant Quillaja saponins (KEECH., et al, 2020). Thus, the definition of the best adjuvant for the composition of a COVID-19 vaccine can be of fundamental importance for both the effectiveness and safety of these new vaccines. So, the data obtained in this work can be of relevant help for the production of safe vaccines

## Declaration of competing interest

Authors have no conflict of interest to declare.

## Acknowledgments

**FAPESP/CEPID/CeTICs, grants nº 2013/07467-1**

**CNPq, grants nº 472744/2012-7**

## Notes

### Competing Interest Statement

The authors have declared no competing interest.

## References

Aguilar, J. C. and Rodriguez, E. G. Vaccine adjuvants revisited. Vaccine, v. 25, n. 19, p. 3752–3762, 2007.

Albertini, R. J.; Anderson, D.; Douglas, G. R.; Hagmar, L.; Hemminki K. K.; Merlo, F. IPCS guidelines for the monitoring of genotoxic effects of carcinogens in humans. Mutat. Res., v. 463, n. 2, p. 111–172, 2000.

Amini, E., Nabiuni, M., Baharara, J., and KAZEM - Parivar Hemolytic and cytotoxic effects of saponin like compounds isolated from Persian Gulf brittle star (Ophiocoma erinaceus). J Coast Life Med, v. 2, n. 10, p. 762–768, 2014.

Antignac, E., Nohynek, G.J, Re, T., Clouzeau, J. and Toutain, T. Safety of botanical ingredients in personal care products/cosmetics. Food and Chemical Toxicology, v. 49, n. 2, p. 324–341, 2011.

Araldi, R.P., Melo, T.C., Diniz, N., Mazzuchelli-De-Souza, J., Carvalho, R. F., BeçAk, W. and Stocco R.C. Bovine papillomavirus clastogenic effect analyzed in comet assay. BioMed Research International, v. 2013, p. 1–7, 2013.

Araldi, R. P., Giovanni, D.N.S., Melo, T.C., Diniz, N. J., Sant’Ana, T.A., Carvalho, R.F., BeçAk, W. and Stocco, R.C. Bovine papillomavirus isolation by ultracentrifugation. Journal of Virological Methods, v. 208, p. 119–24, 2014.

Araldi, R. P., Mazzuchelli-De-Souza, J ., Modolo, D.G., Souza, E.B., Melo, T.C., Spadacci-Morena, D.D., Magnelli, R.F., Carvalho, M.A.C.R., De SáJúNior, P.L. and Carvalho, R.F. Mutagenic Potential of Bos taurus Papillomavirus Type 1 E6 Recombinant Protein: First Description. BioMed research international, v. 2015, 2015.

Audibert, F. Adjuvants for vaccines, a quest. International Immunopharmacology, v. 3, p. 1187–1193, 2003.

Blythe, L.L.; Craig, A.M.; Christensen, J.M.; Appell, L.H.; Slizeski, M.L. Pharmacokinetic disposition of dimethyl sulfoxide administered intravenously to horses. American Journal of Veterinary Research, v. 47, n. 08, p. 1739–1743, 1986.

Bomford, R., Stapleton, M., Winsor, S., Beesley, J.E., Jessup E.A., Price, K.R. and Fenwick, G.R. Adjuvanticity and ISCOM formation by structurally diverse saponins. Vaccine, v. 10, n. 9, p. 572–577, 1992.

Borzacchiello, G and Roperto, F. Bovine papillomaviruses, papillomas and cancer in cattle. Vet. Res. (2008) 39:45

Brayton, C.F. Dimethyl sulfoxide (DMSO): a review. Cornell Veterinarian, v. 76, n. 01, p. 76–90, 1986.

Brito, O.M.; GuimarãEs, M. F. B.; Lanna, C.C.D. Ciclofosfamida e função ovariana. rev bras reumatol, v. 48, n. 1, p. 39–45, 2008.

Campo, M. S.. Vaccination against papillomavirus in cattle. Clinics in dermatology, v. 15, n. 2, p. 275–283, 1997.

Campo, M. S. Papillomavirus research: from natural history to vaccines and beyond. Horizon Scientific Press, 2006.

Chwalek, M., et al. Structure–activity relationships of some hederagenin diglycosides: haemolysis, cytotoxicity and apoptosis induction. Biochimica et Biophysica Acta (BBA)-General Subjects, v. 1760, n. 9, p. 1418–1427, 2006.

Costa, A. F. Lalun, N., Bobichon, H., Plé, K., Voutquenne-Nazabadioko, L. Fármacos com saponósidios. In: Farmacognosia. v.2. ed. 3. Lisboa: Fundação Calouste Gulbenkian, p. 338–357, 2001.

Doorbar, J. The papillomaviruses life cycle. J Clin Virol, v. 32, p.7–15, 2005.

Fabiani, R., Rosignoli, P., De Bartolomeo, A., Fuccelli, R., Servili, M., Montedoro, G., Morozzi, G. Oxidative DNA damage is prevented by extracts of olive oil, hydroxytyrosol, and other olive phenolic compounds in human blood mononuclear cells and HL60 cells. The Journal of Nutrition, v. 138, p. 1411–1416, 2008.

Flores, M.; Yamaguchi, M.U. Teste de micronúcleo: uma triagem para avaliação genotóxica. Revista Saúde e Pesquisa, v.1, n.3, p. 337–340, 2008.

Freitas, A.C., Carvalho C., Brunner., Birgel-Junior, E.H., Dellalibera A.M.M.P. Benesi, F.J., Gregory L., BeçAk, W. and Santos, R.C.S. Viral DNA sequences in peripheral blood and vertical transmission of the virus: a discussion about BPV-1. Braz.J. Microbiol, v.34, p.76–78, 2003.

Friedberg, E. C., Walker, G.C., Siede, W. and Wood, R.D. DNA Repair and Mutagenesis American Society for Microbiology Press. Washington, DC, 2005.

Furesz, J., Fanok, A., Contreras, G. and Becker, B. Tumorigenicity testing of various cell substrates for production of biologicals. Developments in biological standardization, v. 70, p. 233–243, 1988.

Gauthier, C., Legault, J., Girard-Lalancette, K., Mshvildadze, V. and A. Pichette. Haemolytic activity, cytotoxicity and membrane cell permeabilization of semi-synthetic and natural lupane-and oleanane-type saponins. Bioorganic & medicinal chemistry, v. 17, n. 5, p. 2002–2008, 2009.

Glenny, A. T., Pope, C.G., Waddington, H. and Wallace, U. Immunological notes. xvii–xxiv. The Journal of Pathology, v. 29, n. 1, p. 31–40, 1926.

Gupta, R. K.; Relyveld, E. H.; Lindblad, E. B.; Bizzini, B.; Ben-Efraim, S.; Gupta, C. K. Adjuvants-a balance between toxicity and adjuvanticity. Vaccine, v. 11, p. 293–306, 1993.

Gupta, R. K.; Siber, G. R. Adjuvant for human vaccines-current status, problems and future prospects. Vaccine, v. 13, p. 1263–1276, 1995.

Gutiérrez, A.; Rodriguez, I.M.; Jose, C. Chemical composition of lipophilic extractives from sisal (Agave sisalana) fibers. Industrial crops and products, v. 28, n. 1, p. 81–87, 2008.

Heddle, J. A., Salamone, M.F., Hite, M., Kirkhart, B., Mavournin, K., Macgregor, J.G. and Newell, G.W. The induction of micronuclei as a measure of genotoxicity. Mutation. Res. v.123, p.61–118, 1983.

Heuser, V.D., Andrade, V.M., Peres, A., Braga, L.M.G.M. and Chies, J.A.B. Infuence of age and sex on the spontaneous DNA damage detected by micronucleus test and comet assay in mice peripheral blood cells. Cell Biol Int. v.10, p.1223–9, 2008.

Horaud, F. Absence of viral sequences in the WHO-Vero cell bank a collaborative study. Developments in biological standardization, n. 76, p. 43–46, 1992.

Hostettmann, K.; Marston, A. Chemistry and Pharmacology of Natural Products, Saponin. Cambridge University Press, Cftmbridge, 1995.

Keech, C., Albert, G., Cho, I., Robertson, A., Reed, P., Neal, S., Plested, J.S., Zhu, M., Cloney-Clark, S., Zhou, H., Smith, G., Patel, N., Frieman, M.B., Haupt, R.E., Logue, J., Mcgrath, M., Weston, S., Piedra, P.A., Desai, C., Callahan, K., Lewis, M., Price-Abbott, P., Formica, N., Shinde, V., Fries, L., Lickliter, J.D., Griffin, P., Wilkinson, B., AND Glenn, G.M. Phase 1–2 trial of a sars-cov-2 recombinant spike protein nanoparticle vaccine.N Engl J Med 2020; 383:2320–2332

Kensil, C.R., U Patel, D Marciani. Separation and characterization of saponins with adjuvant activity from Quillaja saponaria Molina cortex. The Journal of Immunology, v. 146, n. 2, p. 431–437, 1991.

Kensil, C. R., Wu, J.-Y.; Soltysik, S. Structural and immunological characterization of the vaccine adiuvant QS-21. Pharm Biotechnol. 1995;6:525–41

Kumar, R.; Dwivedi, N.; Singh, R. K.; Kumar, S. Rai, V. P.; Singh, M. A review on molecular characterization of pepper for capsaicin and oleoresin. Internation Jounal of Plant Breeding and Genetics, v.5, 99–110, 2011.

Marigliani, B., Sakauchi, D., Oliveira, H., Sasaki, A., Ferreira Armbruster-Moraes1, E., Mueller, M. and Cianciarullo, A. Intracellular distribuition of recombinant human papilomavirus capsid proteins. Current microscopy contribuitions to advances in Science and technology, v.1, p.678–684, 2012.

Martin, A. R., Martins, M.A., Mattoso, L.H.C. and Silva, O.R.R.F. Caracterização química e estrutural de fibra de sisal da variedade Agave sisalana. Polímeros: Ciência e Tecnologia, v. 19, n. 1, 2009.

Maslov, A. Y.; Vijg, J. Genome instability, cancer and aging. Biochimica et Biophysica Acta (BBA)-General Subjects, v. 1790, n. 10, p. 963–969, 2009.

Modolo,. D.G. Bovine papillomavirus L1 protein (bpv-1): production in bacteria and tobacco plants. 2014. Tese (doutorado). Universidade Estadual de Campinas, São Paulo.

Módolo, D.G., Araldi R.P., Mazzuchelli-De-Souza J., Pereira A., Pimenta D.C, et al. Integrated analysis of recombinant BPV-1 L1 protein for the production of a bovine papillomavirus VLP vaccine. Vaccine, v.35, p. 1590–1593, 2017.

Morein, B., Eriksson, M. V., Sjolander, A.; Bengtsson, K. L. Novel adjuvants and vaccine delivery systems. Veterinary Immunology and Immunopathology, v. 54, p. 373–384, 1996.

Munday, J. Bovine and Human Papillomaviruses: A Comparative Review. Veterinary Pathology, n. June, 2014.

Munday, J. S., Tucker, R.S., Kiupel, M. AND Harvey, C.J. Multiple oral carcinomas associated with a novel papillomavirus in a dog. Journal of Veterinary Diagnostic Investigation, v. 27, n. 2, p. 221–225, 2015. N Engl J Med 2020; 383:2320-2332

Nohynek, J.G., Antignac, E., Re, T. and Toutain, H. Safety assessment of personal care products/cosmetics and their ingredients. Toxicology and applied pharmacology, v. 2, p.239–259, 2010.

O’Hagan, D.T., Singh, M. Microparticles as vaccine adjuvants and delivery systems. Rev.Vaccines, v. 2, p. 269–283, 2003.

O’Hagan, D.T., Lodaya, R.N. and Lofano, G. The continued advance of vaccine adjuvants – ‘we can work it out’. Seminars in Immunology, Volume 50, August 2020, 101426

Ortega, C., FERNáNDEZ-A., Carrillo, J.M. and Romero, P. IL-17-producing CD8+ T lymphocytes from psoriasis skin plaques are cytotoxic effector cells that secrete Th17-related cytokines. Journal of leukocyte biology, v. 86, n. 2, p. 435–443, 2009.

Osada, N., Kohara, A., Yamaji, T., Hirayama, N., Kasai, F., SEKIZUKA Hyperlink Makoto Kuroda, T., AND Hanada, K. The genome landscape of the African green monkey kidney-derived Vero cell line. DNA research, v. 21, n. 6, p. 673–683, 2014.

Peng, S., Wang, J.W., Karanam, B., Wang, C., Huh, W.K.,, Alvarez, R.D., Pai, S.I., Hung, C., Wu, T.C. and Roden, R.B.S. Sequential cisplatin therapy and vaccination with HPV16 E6E7L2 fusion protein in saponin adjuvant GPI-0100 for the treatment of a model HPV16+ cancer. PloS one, v. 10, n. 1, p. e116389, 2015.

Petrovsky, N.; Aguilar, J.C. Vaccine adjuvants: current state and future trends. Immunology and cell biology, v. 82, n. 5, p. 488, 2004.

Ramon, G. Sur l’augmentation anormale de l’antitoxine chez les chevaux producteurs de serum antidiphterique. Bulletin de la Societé Central de Medecine Veterinaire, v. 101, p. 227–234, 1925.

Rand-Luby, L.; Pommier, R.F.; Williams, S.T.; Woltering, E.A.; Small, K. A.; Fletcher, W.S. Improved outcome of surgical flaps treated with topical dimethylsulfoxide. Annals of Surgery, v.224, n.4, p.583–590, 1996.

Reed, S. G., Bertholet, S., Coler, R.N. and Friede, M. New horizons in adjuvants for vaccine development. Trends in immunology, v. 30, n. 1, p. 23–32, 2009.

Ribeiro Filho, J. Cultura do Sisal. Universidade Rural do Estado de Minas Gerais. Viçosa – MG, 1967

Ribeiro-Muller, L.; Muller, M. Prophylactic papillomavirus vaccines. Clinics in dermatology, v. 32, n. 2, p. 235–247, 2014.

Roperto, S., Borzacchiello, G., Esposito, I., Riccardi, M., Urraro, C., LucÀ, R., Corteggio, A., Tatè, R., Cermola, M., Paciello, O. and Roperto, F. . Productive infection of bovine papillomavirus type 2 in the placenta of pregnant cows affected with urinary bladder tumors. PLoS One, v. 7, n. 3, p. e33569, 2012.

Rosenbaum, E.E.; Herschler, R.J.; Jacob, S.W. Dimethyl sulfoxide in musculoskeletal disordens. Journal of the American Medical Association, v.192, n.02, p.309–313, 1965.

Russo, V., Roperto, F., De Biase, D., Cerinop. URRARO C., Munday, J.S. and Roperto, S. Bovine Papillomavirus Type 2 Infection Associated with Papillomatosis of the Amniotic Membrane in Water Buffaloes (Bubalus bubalis). Pathogens 9(4):262, April 2020

Saran, R.; Tiwari, R.K.; Reddy, P.P.; Ahuja, YR. Risk assessment of oral cancer in patients with pre-cancerous states of the oral cavity using micronucleus test and challenge assay. Oral Oncol, v. 44, p. 354–360, 2008.

Schinjs, V.E.J.C. Immunological Concepts of Vaccine Adjuvant Activity. Current Opinion in Immunology, v. 12, p. 456–463, 2000

Sharma, V., Paliwal, R., and Sharma, S. Phytochemical analysis and evaluation of antioxidant activities of hydro-ethanolic extracts of Moringa oleifera Lam. pods. J. Pharm. Res, v. 4, p. 554–557, 2011.

Shope, R.E. Immunization of rabbits to infectious papillomatosis. J Exp Med, v. 65, p.607–624, 1937.

Silva, M.D., Peralba, M.C.R. and Mattos, M.L.T. Determinação de glifosato e ácido aminometilfosfônico em águas superficiais do Arroio Passo do Pilão. Pesticidas: Res.Econotoxicologia Meio Ambiente, n.13, p.19–28, 2005.

Silva, M. A R., De Albuquerque, B.M.F., Pontes, N.E., Coutinho, L.C.A., LeitãO, M.C.G., Reis, Castro, R.S. and Freitas, A.C. Detection and expression of bovine papillomavirus in blood of healthy and papillomatosis-affected cattle. Genetics and molecular research: GMR, v. 12, n. 3, p. 3150–6, 2013.

Singh N.P., Mccoy M.T., Tice R.R., Schneider E.L. A simple technique for quantitation of low levels of DNA damage in individual cells. Exp Cell Res, v. 175, p.184–191, 1988.

Singh, M.; O’Hagan, D. T. Recent advances in veterinary vaccine adjuvants. International Journal of Parasitology, v. 33, p. 469–478, 2003.

Soltani, M., Parivar, K., Baharara, J., Kerachian, M.A., and Asili, J. Hemolytic and cytotoxic properties of saponin purified from Holothuria leucospilota sea cucumber. Reports of biochemistry & molecular biology, v.3, n. 1, p. 43–50, 2014.

Soltysik, S., Wu, J.Y., Recchia, J., Wheeler, D.A, Newman, M.J., Coughlin, R.T and Kensil, C.R. . Structure/function studies of QS-21 adjuvant: assessment of triterpene aldehyde and glucuronic acid roles in adjuvant function. Vaccine, v. 13, n. 15, p. 1403–1410, 1995.

Stocco Dos Santos, R. C., Lindsey, C.J., Ferraz, O.P, Pinto, J.R, Mirandola, R.S, Benesi, F.J, Birgel, E.H, Pereira, C.A and BeçAk, W. Bovine papillomavirus transmission and chromosomal aberrations: an experimental model. The Journal of general virology, v. 79 (Pt 9), p. 2127–35, 1998.

Sun, H.X., Xie, Y., Ye, Y.P. Advances in saponin-based adjuvants. Vaccine 27:1787–1796, 2009b.

Tice, R. R., Agurell, E., Anderson, D., Burlinson, B., Hartmann, A., Kobayashi, H., Miyamae, Y., Rojas, E., Ryu, J.C. and Sasaki, Y.F. Single cell gel/comet assay: guidelines for in vitro and in vivo genetic toxicology testing. Environmental and molecular mutagenesis, v. 35, n. 3, p. 206–221, 2000.

Vincent-Falquet, J. C., Peyron L., Souvras M., Moulin J.C., Tektoff J. and Patet J Qualification of working cell banks for the Vero cell line to produce licensed human vaccines. Developments in biological standardization, v. 70, p. 153–156, 1988.

Voutquenne, L., Lavaud, C., Massiot, G. and Le Men-Olivier, L. Structure-activity relationships of haemolytic saponins. Pharmaceutical biology, v. 40, n. 4, p.253–262, 2002.

Wang, Y., Zhang, Y., Zhu, Z., Zhu, S., Li, Y., Li, M. and Yu, B. Exploration of the correlation between the structure, hemolytic activity, and cytotoxicity of steroid saponins. Bioorganic & Medicinal Chemistry. v. 15, p .2528–2532, 2007.

Wang, J. W., Roden, R. B.S. L2, the minor capsid protein of papillomavirus. Virology, v. 445, n. 1, p. 175–186, 2013.

Yaguiu, A., Dagli, M.L.Z., Birgel Jr, E.H., Reis, B.C.A.A., Ferraz, O.P., Pituco, E.M., Freitas, A.C., BeçAk, W. and Stocco, R.C. Simultaneous presence of bovine papillomavirus and bovine leukemia virus in different bovine tissues: in situ hybridization and cytogenetic analysis. Genet.Mol.Res. v.7, p.487–497, 2008.

